# Functions of *C. elegans* neurons from synaptic connectivity

**DOI:** 10.1101/2024.03.08.584145

**Authors:** Scott W. Emmons

## Abstract

Despite decades of research on the *C. elegans* nervous system based on an anatomical description of synaptic connectivity, the circuits underlying behavior remain incompletely described and the functions of many neurons are still unknown. Updated and more complete chemical and gap junction connectomes of both adult sexes covering the entire animal including the muscle end organ have become available recently. Here these are analyzed to gain insight into the overall structure of the connectivity network and to suggest functions of individual neuron classes. Modularity analysis divides the connectome graph into ten communities that can be correlated with broad categories of behavior. A significant role of the body wall musculature end organ is emphasized as both a site of significant information convergence and as a source of sensory input in a feedback loop. Convergence of pathways for multisensory integration occurs throughout the network — most interneurons have similar indegrees and outdegrees and hence disperse information as much as they aggregate it. New insights include description of a set of high degree interneurons connected by many gap junctions running through the ventral cord that may represent a previously unrecognized locus of information processing. There is an apparent mechanosensory and proprioceptive field covering the entire body formed by connectivity of the many mechanosensory neurons of multiple types to two interneurons with output connections across the nervous system. Several additional significant, previously unrecognized circuits and pathways are uncovered, some involving unstudied neurons. The insights are valuable for guiding theoretical investigation of network properties as well as experimental studies of the functions of individual neurons, groups of neurons, and circuits.

## Introduction

The nervous system evaluates and integrates sensory information of various kinds from the external environment and from internal sensors to generate coherent, adaptive outputs — behavioral, physiological, reproductive. The functions of individual neurons or brain regions in this process has been a longtime focus of neuroscience research. These functions are determined by the roles each neuron plays in the network of cellular communications created by synaptic and extrasynaptic signaling. A description of the set of chemical and electrical synapses visible by electron microscopy has been available for the *C. elegans* nervous system for almost 40 years (3), while studies of the extrasynaptic neuropeptidergic signaling network have more recently begun to emerge (4, 5).

The anatomical synaptic network of the *C. elegans* nervous system has recently been reanalyzed and extended across the entire animal for both adult sexes and to larval stages (2, 6). From their physical connectivity and structures, together with results of many experimental studies, it has been possible to assign functions to many of the neurons. Sensory neurons often have identifiable dendritic endings in sensory structures while motor neurons have output onto muscles. However, almost all *C. elegans* neurons are multifunctional, being both pre- and post-synaptic to other neurons as well as having sensory properties and making neuromuscular junctions. Moreover, considerable cross connectivity, especially among interneurons, creating a complex network, has made the interpretation of functions of many neurons problematic. At present, the functions of 57% (26/46) of the neuron classes identified as interneurons (81 neurons total) remain largely unknown (see WormAtlas.org). Uncertainty is unevenly distributed—certain regions of the nervous system are better known than others. For example, the availability of behavioral assays for chemotaxis and social behaviors have resulted in elucidation of circuits involved in navigation in the chemical environment and responses to other nematodes. By contrast, the circuits that process information from sensors facing into the pseudocoelom, for example, are less well known.

The present work takes advantage of the new reconstructions of the chemical and gap junction synaptic connectomes of the adults of both sexes to extend the assignment of functions to neurons (**Supplementary File 1)** (2). From the analysis, it is possible to suggest some functions for most of the previously less well-known interneurons. Several neurons and sets of neurons emerge with unexpected significance. A network of high degree interneurons that extend across the animal in the ventral cord and connect heavily through gap junctions are connected to each other and collectively reach the entire nervous system in a small number of synaptic steps. It is suggested that this central network may represent a previously undescribed locus of integration. The pattern of convergence and divergence of the connectivity from sensory inputs to muscle outputs emphasizes the extent to which the process of multisensory integration occurs throughout the network and involves all cell types at all levels, including the muscle cells themselves.

## Results

### The *C. elegans* nervous system has a modular architecture

As a first step towards assigning functions to neurons, the neurons and muscles may be partitioned into subgroups by their connectivity. The *C. elegans* connectome has been subjected to graph analysis previously (8–12). Various graph theoretic algorithms are available for identifying subgroups (modules or communities) where the probability of an edge between members of communities is significantly greater than expected if edges are distributed randomly. In such community detection, the boundaries in optimal partitions vary by algorithm and by the value of an unavoidable arbitrary threshold parameter, as generally it is not possible to reach a unique solution. Here, as a starting point for analysis, the spectral method of Leicht and Newman (7) is used. This algorithm partitioned the connectivity graph of the adult hermaphrodite of Cook et al. (2) into 10 communities (**Figure 1**). The graph used is a weighted graph where the values of the edge weights are the number of EM serial sections of connectivity summed over the often-multiple synapses connecting pairs of neurons, neurons and muscles, and muscle cells. Chemical and gap junctional graphs, the latter treated as two opposing directed edges with edge weights divided by ½, were added together to create a single graph with 473 nodes and 6951 edges (**Supplementary File 2)**. (Communities created for the chemical and gap junction graphs separately are given in **Supplementary File 3 and 4**).

**Figure 1.**
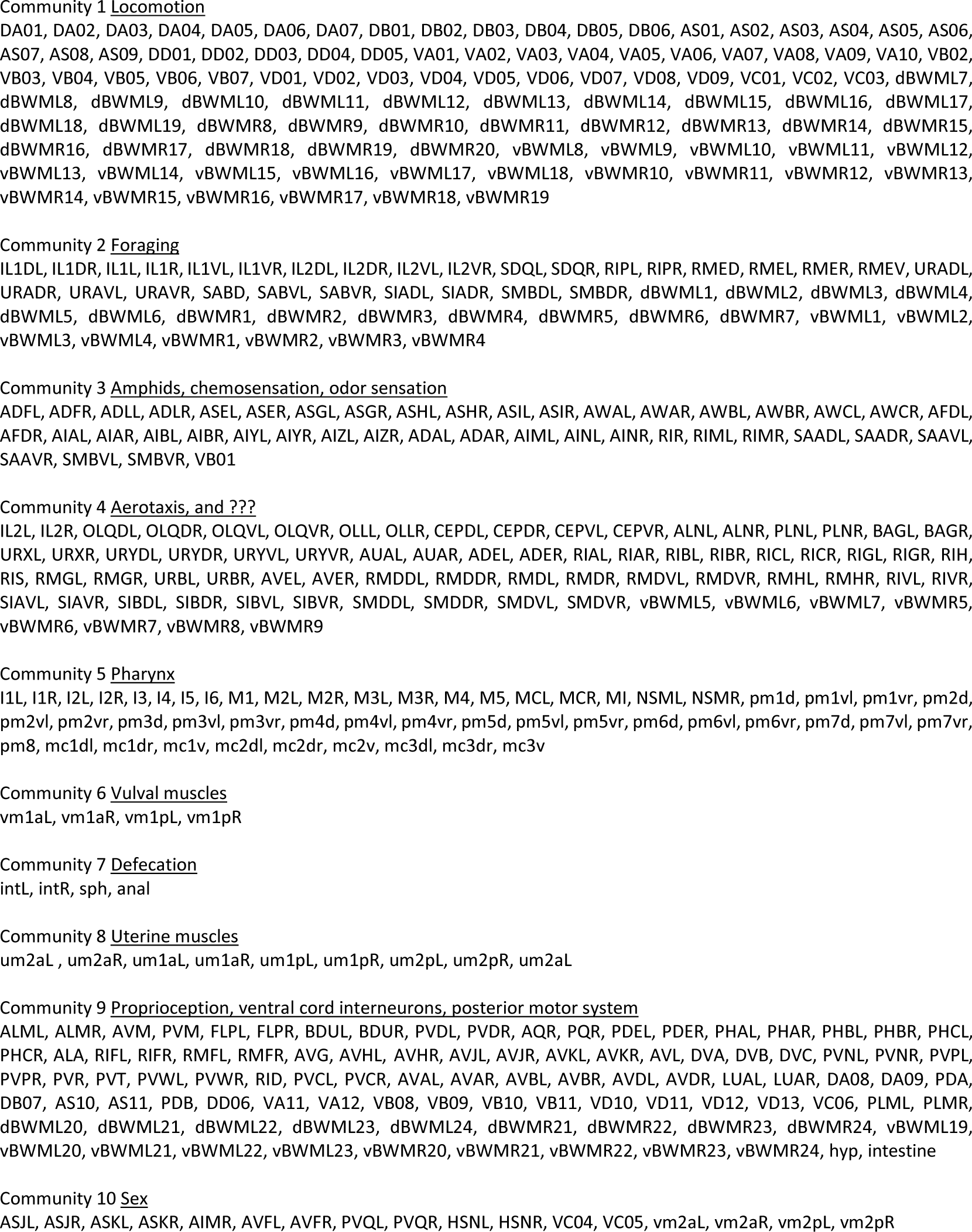
Assignment of the neurons and muscles of the adult hermaphrodite to ten communities by the spectral method of Leicht and Newman (7). The connectivity matrix used, based on the data of Cook et al. (2) (**Supplementary File 1**), was the sum of the weighted chemical matrix plus the weighted, symmetrical gap junction matrix with gap junction weights divided by 2 (since each edge is counted twice, as directed edges in each direction) (**Supplementary File 2**). Modularity parameter Q = 0.517. P < 10^-6^

The division into communities makes clear distinctions among various types of inputs and outputs. Community 1 contains most, but not all, of the longitudinal body wall muscles and the excitatory and inhibitory ventral cord motor neurons that innervate them. Communities 6, 7, and 8 separate out additional small muscle groups involved in egg laying and defecation. Community 5 isolates all the muscles and neurons of the pharynx and is not considered further here. The remaining four communities represent vertically structured silos of information flow from sensory neurons of a particular type all the way to motor neurons and even in some cases specific body wall muscles.

It is important to emphasize that description of the network in this manner is artificial in requiring nodes to be placed into one or another of a small number of groups. A 2D, spring-electric or force directed layout of the *C. elegans* nervous system that, like modularity analysis, arranges nodes according to the amount of connectivity between them, is given as **Supplementary File 5** (2). While it groups neurons and muscles consistent with the modularity analysis, it shows no obvious boundaries or separations between many of the modules. The precise location of the boundary drawn by the modularity algorithm may be quite arbitrary and highly sensitive to particular connections. Thus, as an example, the separation of SIADL and SIADR in comm 2 away from SIAVL, SIAVR, and four SIB neurons in comm 4 is not reflected in the 2D layout and probably does not indicate a difference in function.

### Convergence onto the muscles and motor neurons

In the overall process of multisensory integration, information from multiple sensory streams is brought together for control of effector output. This process can be dissected stepwise in terms of convergence onto individual neurons and muscles—the number of neurons targeting each cell, its *indegree*. By this measure, in the *C. elegans* nervous system muscle cells are important sites of convergence. *C. elegans* locomotion and posture is determined by the four rows of longitudinal body wall muscles (95 mononucleate muscle cells) (**Figure 2A**). Neurons of all types, sensory neurons and interneurons as well as neurons classified as motor neurons, make neuromuscular junctions. The majority of muscle input (the sum of nmj edge weights in the chemical graph) (91%) is from three major motor neuron classes: *head motor neurons* (14%), *sublateral motor neurons* (41%), and *ventral cord motor neurons* (37%).

**Figure 2.**
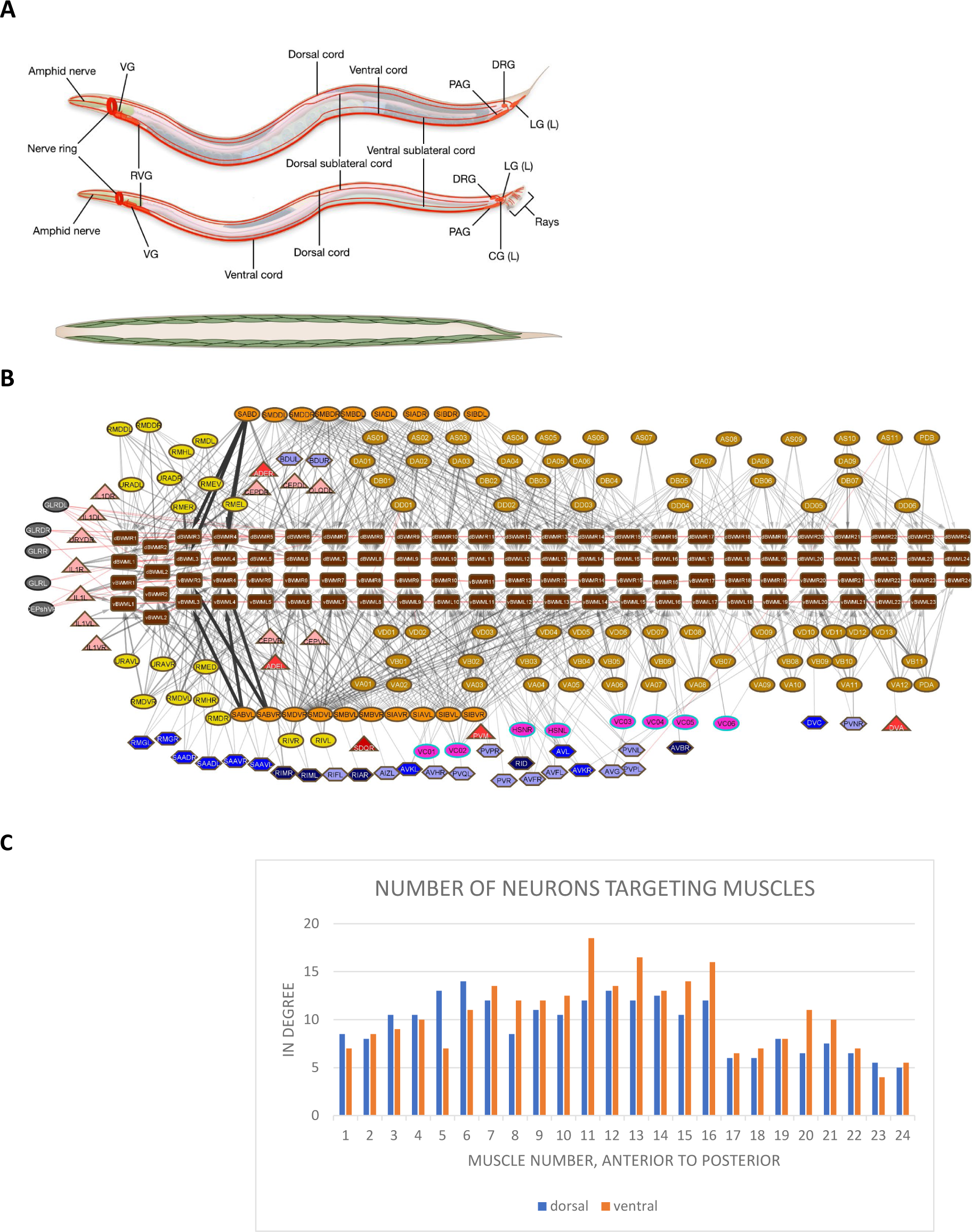
**A.** The nervous system and body wall muscles of *C. elegans* (*upper*: hermaphrodite; *lower*: male). The major ining the cell bodies, and the primary process tracts are shown. Neurons are largely unbranched; processes s with a small number of neighbors to which they make *en passant* synaptic connections. CG, cloacal G, dorsorectal ganglion; LG, lumbar ganglion; PAG, pre-anal ganglion; RVG, retrovesicular ganglion; VG, on. The 95 body wall muscles are arrayed in four quadrants: dorsal left, dorsal right, ventral left, ventral neighbor connections of the longitudinal body wall muscles (brown rectangles). Neurons classified as motor ovals. For interpretation of other node shapes and colors, see legend to **Figure 3** (2). Black lines with chemical connections; red lines: gap junction connections. **C.** Number of neurons targeting the body wall ue shown is the average of the left/right pairs at each longitudinal and dorsoventral position.

**Figure 2B** shows how input from all the various classes of neurons making neuromuscular junctions is distributed across the dorsal and ventral longitudinal muscle chains. On average, each muscle cell receives chemical input from 10 neurons (9.5 in the dorsal set, range 5-12.5, left/right averaged; 10.5 in the ventral set, 4-18.5) **(Figure 2C)**. Each muscle cell also has gap junctions with its neighbors. Thus, muscle cells are strongly convergent.

This input to the muscles comes from all the communities. Each community contains a different subset of the three major motor neuron classes—most ventral cord motor neurons in comm 1, sublateral motor neurons in comm 3, head motor neurons and sublateral motor neurons in comm 4, additional ventral cord motor neurons in comm 9. In addition, IL1 sensory motor neurons, in comm 2, and VC hermaphrodite specific motor neurons, in comm 10, provide additional significant input (**Figure 3**).

**Figure 3.**
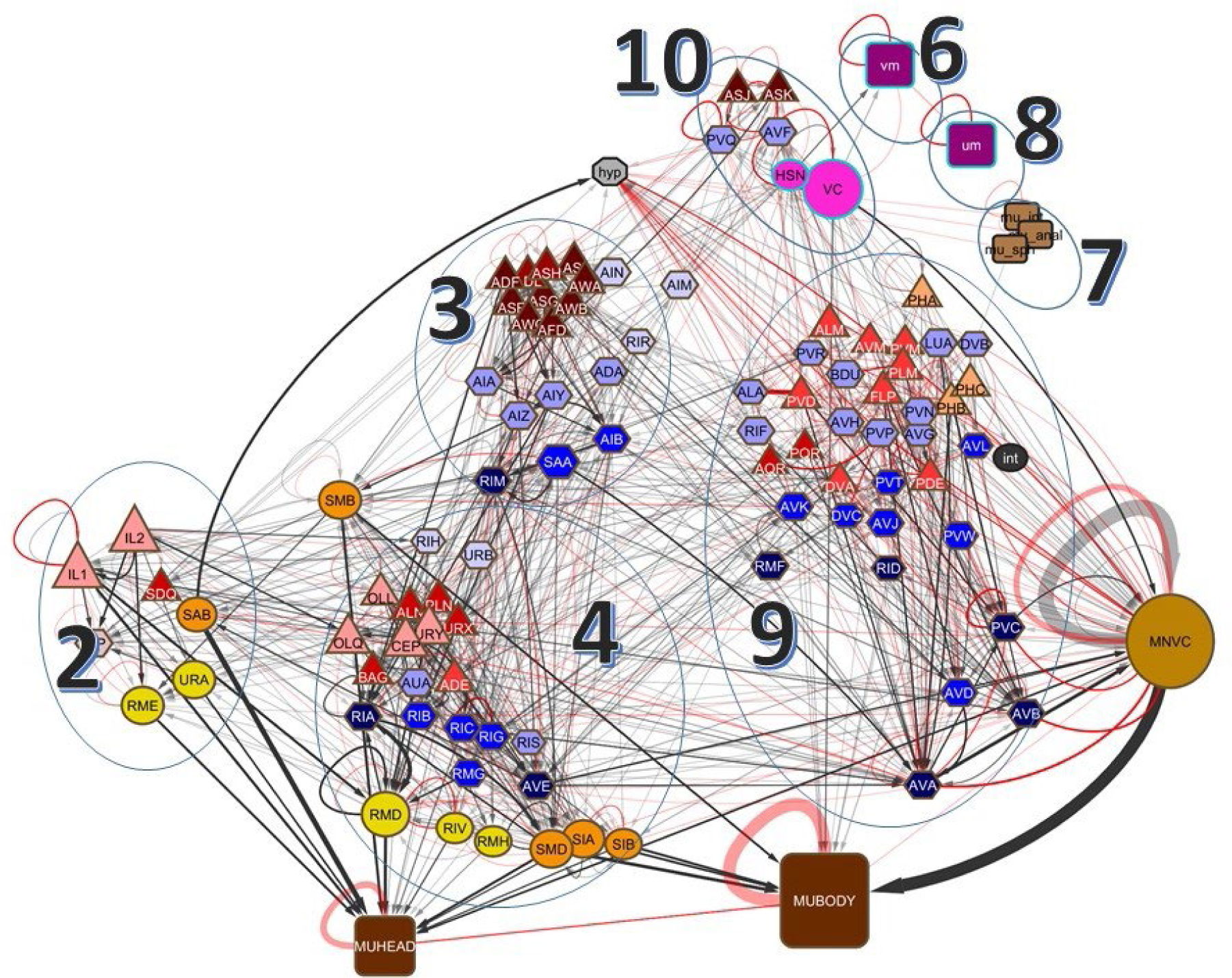
Hierarchical arrangement of the hermaphrodite connectome with neurons clustered by mmunity (not shown: comm 1, containing many motor neurons and bodywall muscles; comm 5, pharyngeal neurons and muscles, connected via RIP). The neurons and muscles are grouped by class (**Supplementary File 9**); node sizes roughly represent the number of cells in each class. Members of several classes lie in different communities: SMBDL and SMBDR are in comm 2, SMBVL d SMBVR are in comm 3; AIML is in comm 3, AIMR is in comm 10; vental cord motor neurons NVC) are in comm 1 and comm 9, bodywall muscles (MUHEAD, MUBODY) are in comm 1, comm comm 4, and comm 9. This is a direct adaptation of **Figure 2** of Cook et al. (2); the symbols and ors are those of that figure. *Shapes*: triangles, sensory neurons; hexagons, interneurons; circles, tor neurons; squares, muscles. Colors: dark red, amphid neurons; medium red, oxygen sensors; ht red, proprioceptors; pink, cephalic sensory neurons; yellow, head motor neurons; orange, blateral motor neurons; tan, ventral cord motor neurons; bright and dark pink nodes with quoise borders are hermaphrodite-specific sex neurons. For the interneurons, the four blue ors, lightest to darkest, represent the four interneuron layers assigned in Cook et al (2). *hyp* is hypodermis. Black lines with arrowheads: chemical connections; red lines: gap junction nnections; line thickness proportional to connection weight, based on number of synapses and apse sizes.

The motor neurons themselves are strongly convergent; input comes from both sensory neurons and interneurons, as well as from other motor neurons (**Figure 4**). The average indegree for the three major motor neuron classes in the combined chemical plus gap junction graph (**Supplementary File 2)** are 19.4 for the head motor neurons, 20.1 for the sublateral motor neurons, and 13.7 for the ventral cord motor neurons. Much of the input to the ventral cord motor neurons (62% of the weighted chemical input and 55% of the weighted electrical input) comes through the so-called command interneurons, interneurons running through the ventral chord that have inputs across the entire chain of ventral cord motor neurons (AVA, AVB, AVD, and PVC). The average indegree of the command interneurons as a group in the combined chemical and gap junction graph is 48.5 (excluding the gap junction connections to the ventral cord motor neurons, which are most likely best considered output). Command interneurons are members of a so-called rich club of high degree, interconnected neurons (10, 13). However, calcium imaging of brainwide activity patterns confirms that the command interneurons are only one element of a motor command network dispersed widely through the nervous system (13, 14).

**Figure 4.**
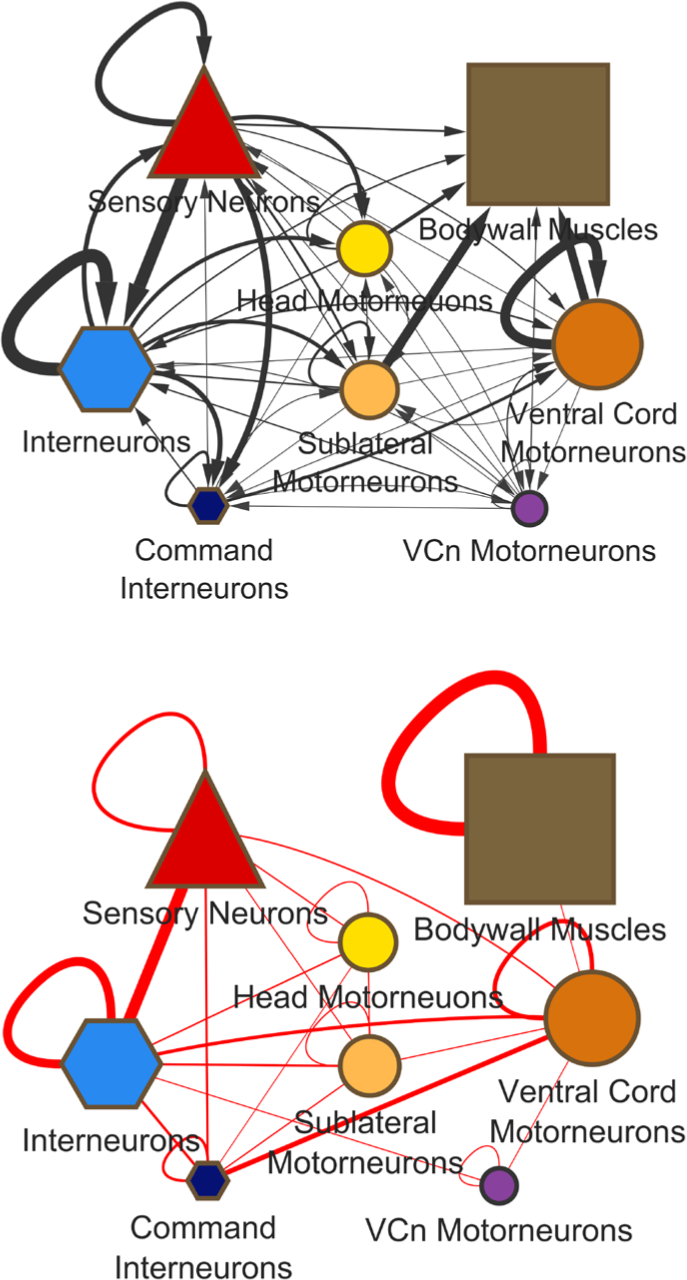
Input to the motor system comes from all cell types. Much of the information progressing through the network converges at the level of the motor neurons. Symbols and colors as for **Figure 2**.

### Sensory and Behavioral Functions of the Modules

Each of the four large sensory modules appears to bring together a spectrum of similar or related sensory inputs that have a common implication for behavioral output. Groupings corresponding roughly to communities 2, 3, and 4 were pointed out by White et al (3) (**Figure 21a, b, c**). The sensory neurons in comm 2 are a subset of those neurons having sensory endings in the inner labial sensilla, IL1 and IL2, and also include the SDQ neurons, which lie along the body and apart from participation in oxygen sensation are of no known function. IL1 and IL2 assess chemical and tactile information near or at the nose for positioning the nose in foraging (15); both target the body wall muscles in the head via head motor neurons RME and URA, while IL1 targets these muscles directly as well. SDQR targets the RMH head motor neurons.

The sensory neurons in comm 3 have endings in the amphid sensilla. All the amphid neurons are in this module except for pheromone sensors ASJ and ASK, which are in comm 10. Amphid neurons detect soluble and volatile chemicals in the environment, as well as temperature, factors that may exist in long-range gradients. Such gradients provide a global coordinate system for the worm to use for navigational guidance, for example to navigate to a point where it last obtained food. Here the relevant behavioral output is the durations of the bouts of forward and backwards locomotion and the deep bends and turns that punctuate them, known as pirouettes. The major interneuron targets of amphid sensory neurons, AIA, AIY, and AIZ, are in layer 3 of the hierarchical network layout (**Figure 3**). From these, pathways of connectivity lead to the head motor neurons and to the ventral cord motor neurons via the command interneurons (16).

The sensory neurons of comm 4, like those of comm 2, have both chemical and mechanosensory endings in the sensilla of the nose, and also include several with processes lying along the body or facing the coelomic cavity. The major interneuron targets in this module, RIB, RIC, RIG, and RMG, are in layer 2 (**Figure 3**). This module includes the so-called “neck” muscles, ventral body wall muscles 5-9, as well as the head motor neurons RMD, RMH, and RIV, which target this muscle group and receive inputs from RIC, RIG, and RMG. The AVE interneuron controls both dorsal and ventral ventral chord motor neurons 1-4. Modalities of the sensors of comm 4 are not well characterized but include oxygen sensation. The selective innervation of muscles in the head and neck region suggests control of searching activity that involves movement of the head.

A possible generalization emerges from this partitioning of the network. Each sensory module may control a separate searching strategy: comm 2 for targets right at the nose (foraging), comm 4 for targets or goals nearby (possibly behavior known as steering), and comm 3 for locating targets or goals farther away (runs punctuated by pirouettes). Such strategies might respectively involve regulating movement of the nose, the head, and the entire body. This viewpoint helps to explain the many pathways to the motor system with independent contributions from each of the sensory modules (**Figure 3**).

In contrast to communities, 2, 3, and 4, which include sensation of signals in the external environment, comm 9 appears to assess information on the condition of the body itself. It includes the various types of mechanoreceptors (touch neurons, deirids, PVD and FLP) as well as neurons with sensory endings facing into the body cavity (AQR and PQR). Finally, module 10 brings in information relevant to reproductive behavior. The sex-specific circuits involving AVF and PVQ are shown in Cook et al. (2), **Figure 6g,h**.

### Integration of information higher in the network

It might be expected that sensory information with a common implication for behavior would first be compared and conflicts over priority resolved within each module, and then this unified modular output would be brought together with that of the other modules for prioritization and regulation of the motor neurons and muscles. Such integration would be a major function of the interneurons. Some would be involved in processing inputs within modules, while others would be nodes for comparing modular outputs. In this scheme, interneurons would in general have an overall convergent function, that is, they would have higher indegrees than outdegrees, reducing a larger number of incoming information streams to a smaller number of outputs. Interneuron connectivity indicates the above scheme is too simplistic. For most interneurons, the ratio of indegree to outdegree is close to 1 (**Figure 5**). That is, they do not have an overall convergent function. In each module there are some interneurons that in fact disperse information more than they aggregate it. Notably, as an example, the two interneurons in comm 10, AVF and PVQ, which receive sexually relevant inputs, disperse this information widely across the nervous system. The most strongly convergent interneurons are pre-motor interneurons, that is, neurons with a preponderance of output onto motor neurons. In addition to the command interneurons discussed above, these include RIA and AVE. (AVE has been considered a command interneuron heretofore, even though it only synapses onto the anterior subset of ventral cord motor neurons.)

**Figure 5.**
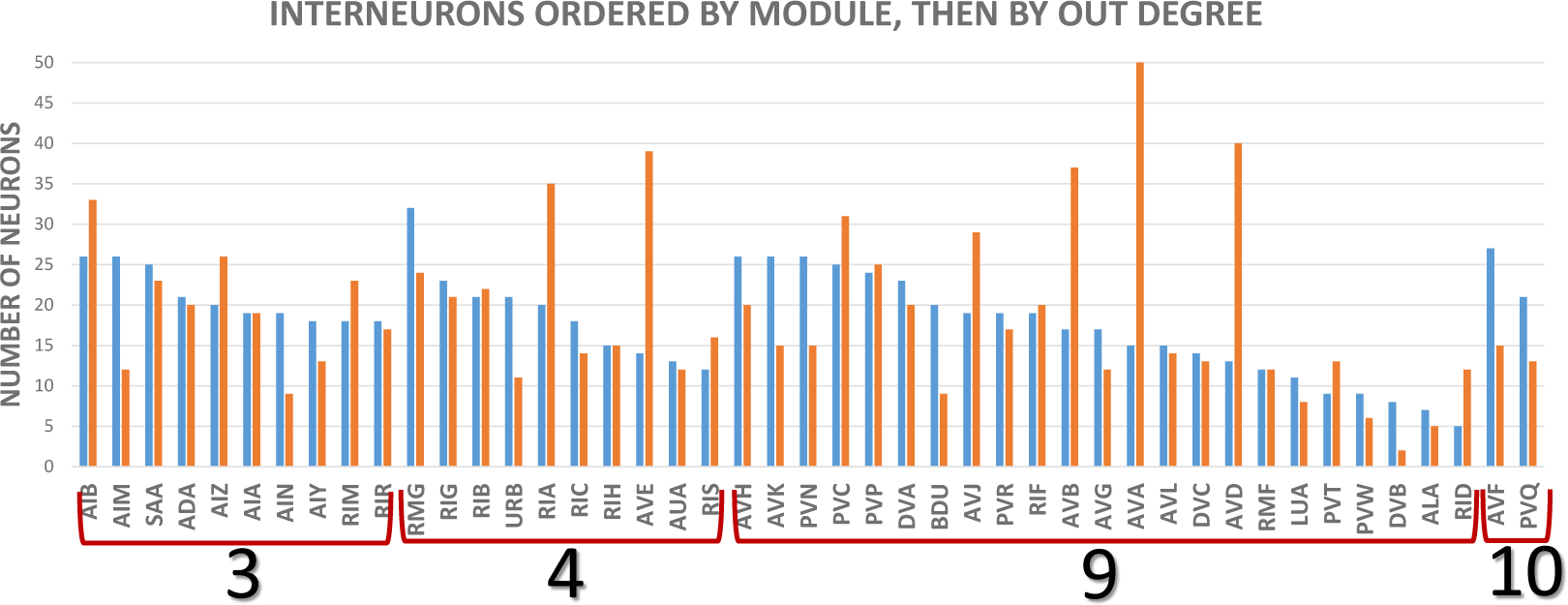
Out (blue) and indegrees (orange) for classes of interneurons. These are the degrees of classes onto other classes, taken from the cell class adjacency matrix of **Supplementary File 9;** ventral cord motor neurons, muscles in the head, and remaining muscles through the body, are counted as one node. (DVA has been classified as a sensory neuron previously because it is stretch sensitive (1).

### A central role for ventral cord interneurons

The high level of connectivity of the interneurons, particularly their generally high chemical outdegrees and large amount of electrical connectivity, which is of uncertain directionality, is the reason interneuron functions have been difficult to parse out from connectivity **(Figure 6)**. The interneurons with the highest degrees, particularly gap junction degrees, include interneurons that run across the body in the ventral cord between the nerve ring and the tail ganglia. All are in community 9 except AVF and PVQ, which are in community 10. This group includes the well-known command interneurons AVA, AVB, AVD, and PVC, but the remainder are little studied. Collectively, the 13 classes of non-command, ventral cord interneurons synapse onto a large fraction of all the neurons in the nervous system — by chemical connections, 59% in one step, 98% in two steps; by gap junctions, 41% in one step, 85% in two steps. Moreover, they are heavily connected to each other **(Figure 7)**.

**Figure 6.**
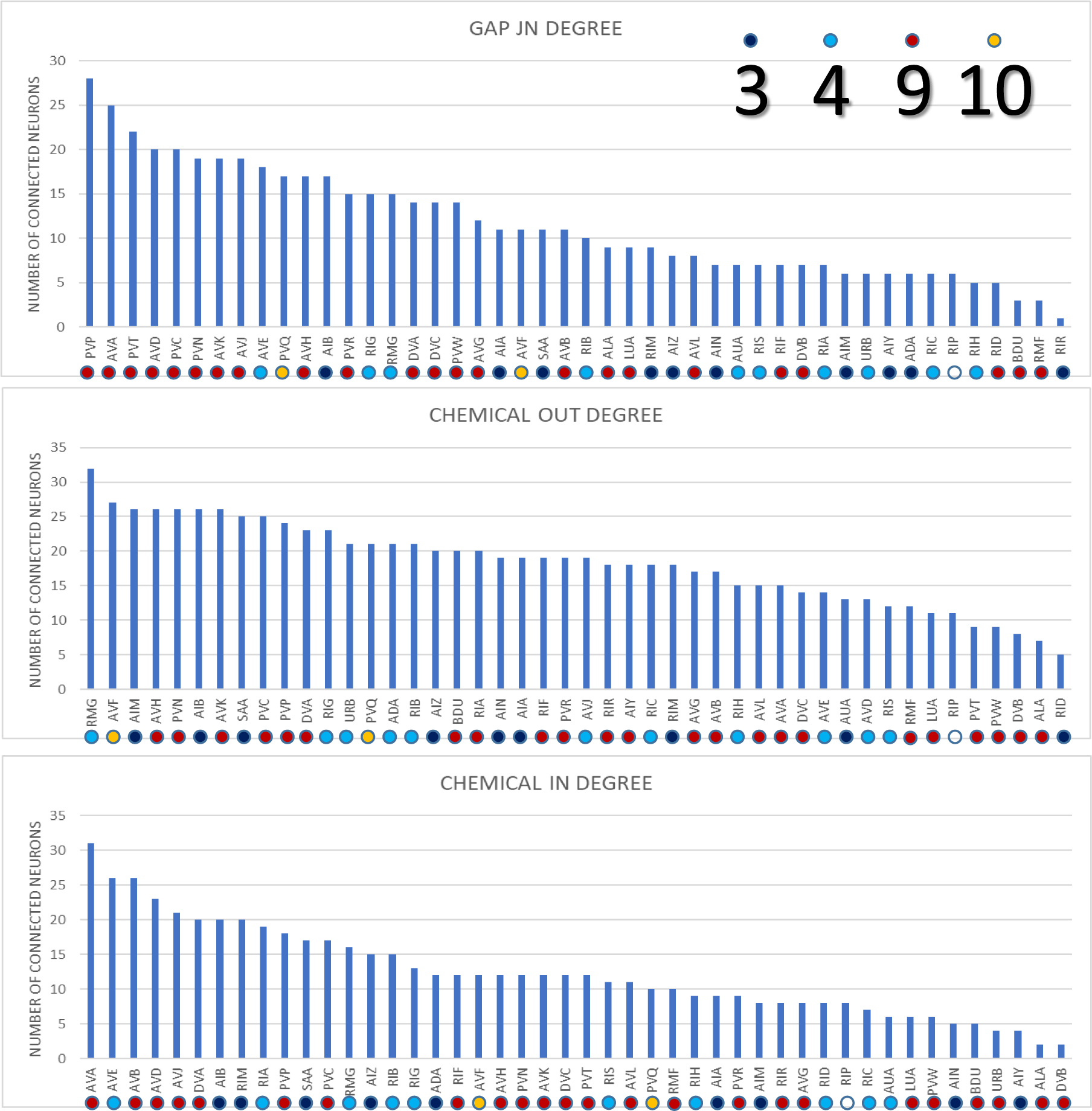
Degrees of interneurons; modules shown by color code. From cell class adjacency atrix **Supplementary File 9**, as in **Figure 5**.

**Figure 7.**
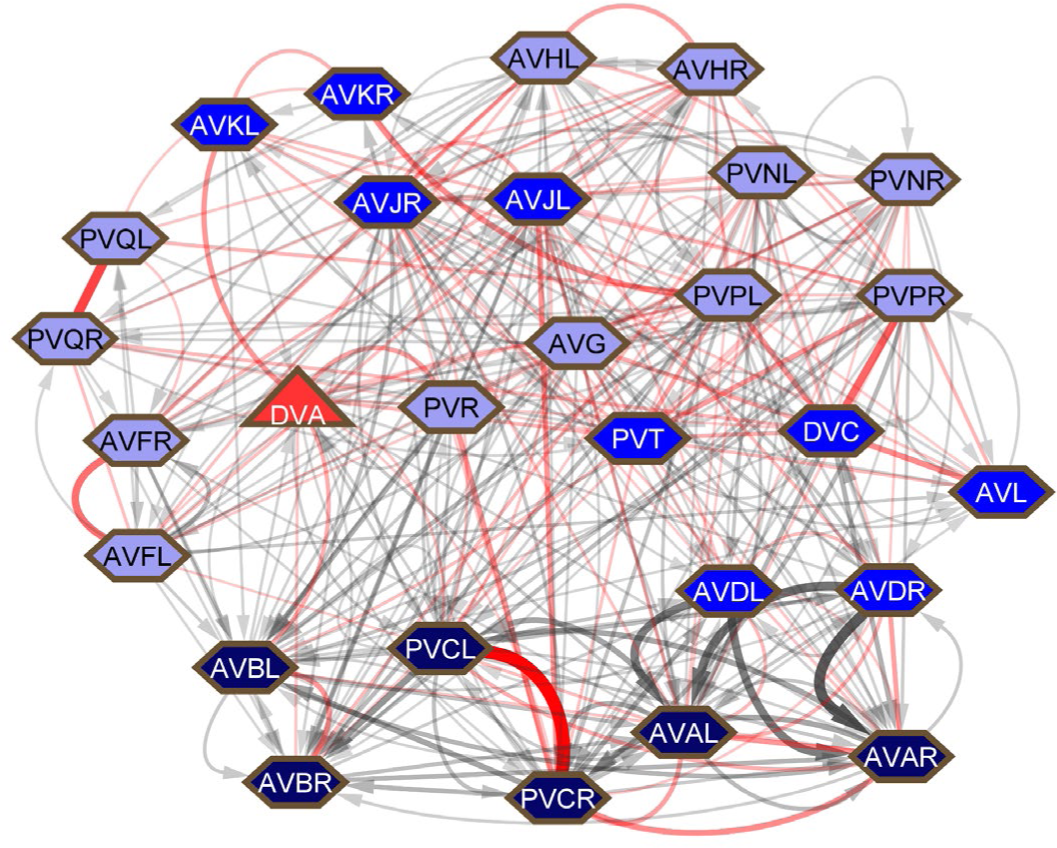
Connections among ventral cord interneurons. s and colors as for **Figure 2**.

This would appear to be an important central network for integrating and dispersing information across the nervous system. In the whole-animal somatic nervous system connectivity graphs of Cook et al. (2), there are a total of 4167 chemical plus gap junction synapses in the nerve ring, usually considered the central locus of nervous system integration, and 4897 (54% of the total) outside the nerve ring in the ventral cord and tail (**Supplementary File 6**, updated from SI3, Synapse Lists of Cook et al (2)). **Figure 8** shows how the inputs and outputs of the non-command, ventral cord interneurons are distributed across the body. While some, like DVA, PVR, and PVT, appear to bring inputs from outside the nerve ring into it, others have inputs and outputs more uniformly distributed. There are a remarkable number of gap junctions in the tail. (The possibility that this previously unnoted imbalance is a reconstruction artifact is reviewed in the Discussion.)

**Figure 8.**
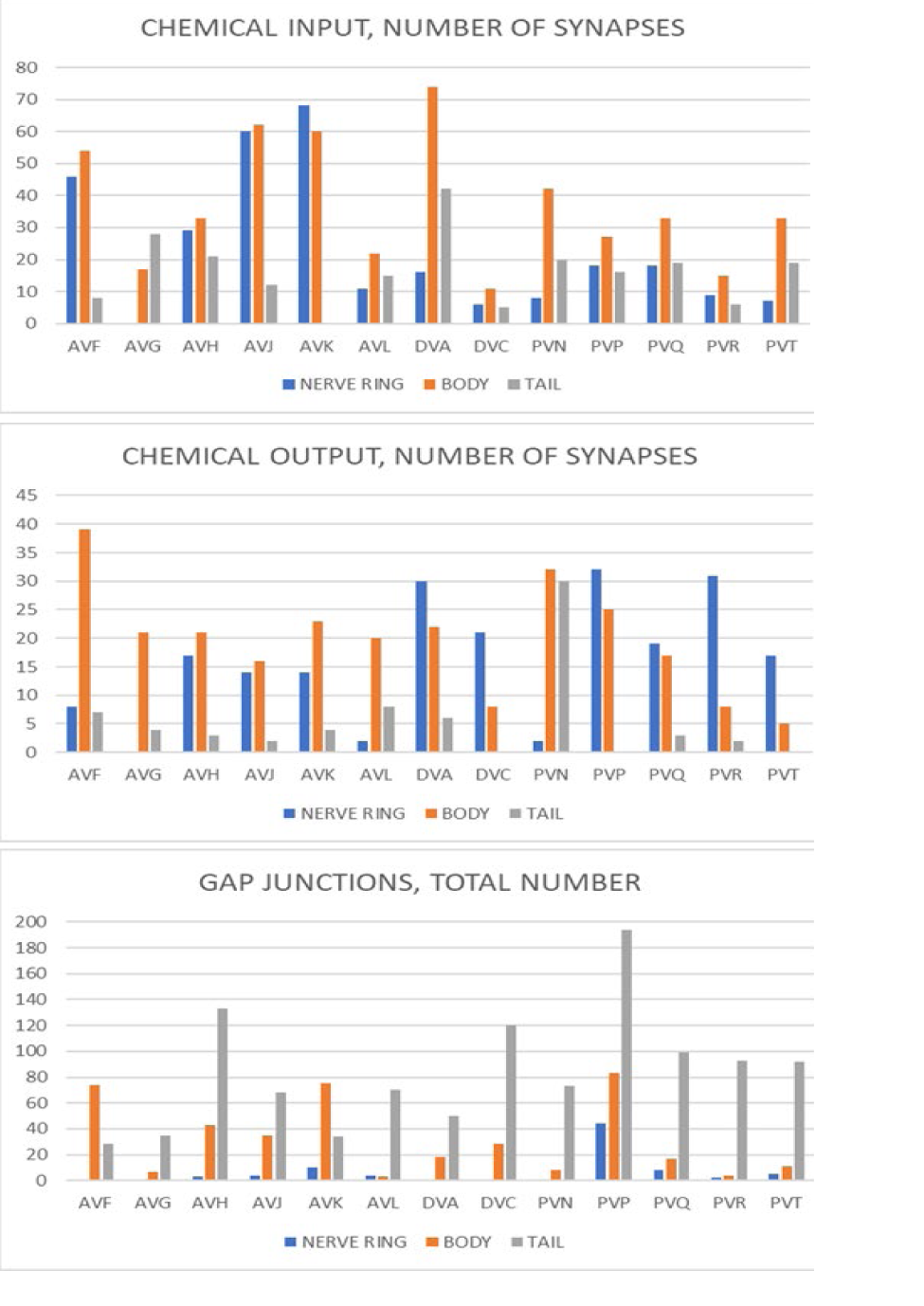
Distribution of synapses by body region along ventral cord interneurons. From **Supplementary File 6**. The body region here includes the ventral ganglion, the retrovesicular ganglion, and the ventral cord. The tail region includes the pre-anal ganglion, the dorsorectal ganglion, and the lumbar ganglia (See **Figure 2A**).

Two measures of network topology are *betweenness centrality*, a node property that reflects the number of shortest paths between pairs of nodes that pass through it, and *rich-club coefficient*, a measure that identifies high degree neurons that are strongly connected to each other. Nine of the top 20 interneurons by betweenness centrality in the combined hermaphrodite chemical plus gap junction graph are members of comm 9 (5 are command interneurons, 4 are non-command ventral cord interneurons), 6 are in comm 3, and 5 are in comm 4 (**Supplementary File 7**). Among 21 neurons identified as rich clubs in the combined chemical plus gap junction graph, 11 are in comm 9 (8 command interneurons, 3 non-command ventral cord interneurons), 3 are in comm 3, 6 are in comm 4, and 1 is in comm 10) (13).

Further emphasizing their important role in nervous system integration, not only are the non-command ventral cord interneurons in comm 9 heavily connected by synapses, several are also the highest degree neurons in the extra-synaptic peptidergic communication network: AVK, PVT, PVQ, DVA, and PVR — all are higher than the next highest neurons, which are the command interneurons AVA and PVC (5).

### Sensory networks for the conditions of the body

Whereas the many sensory neurons for assessing external environmental conditions are grouped in comm 2, 3, and 4 and largely have sensory dendrites in the head and outputs focused on interneurons in the nerve ring, the various types of mechanosensory neurons with dendrites distributed across the body are in comm 9 and have important interneuron targets in this community. Two of these interneurons, DVA and PVR, receive inputs from the entire spectrum of mechanosensors (**Figure 9**). In the hermaphrodite reconstruction (but not the male), PVR and DVA are joined by 8 gap junctions and additional reciprocal chemical connections. They share output to command interneurons AVB and PVC, but otherwise their output connections are distinctive. DVA has output onto AVA and ring interneurons involved in the navigation circuitry, while PVR apparently regulates pharyngeal function through output onto RIP, the interneuron that connects to the pharyngeal nervous system, and IL1, which also targets RIP. As noted above, not only are DVA and PVR high degree neurons in the synaptic network (**Figure 6**), they are among the highest degree neurons in the peptidergic, extrasynaptic communication network. DVA and PVR themselves appear to have sensory function. DVA is stretch sensitive and PVR has a process extending into the tail whip that sometimes contains a cilium (1, 17). Nevertheless, these neurons are perhaps best viewed as interneurons receiving input from a somatosensory receptive field that consists of the entire body — surface, cuticle, and coelomic cavity. In addition to output onto navigational interneurons, DVA has output onto sublateral motor neurons whose activity may be relevant to bodywide muscle tone. Such tone, as well as pharyngeal pumping, must impact pressure in the body, which needs to be kept sufficiently high for function of the cuticular exoskeleton but not so high as would cause the cuticle to burst. It has been reported that touch can stop pharyngeal pumping (18).

**Figure 9.**
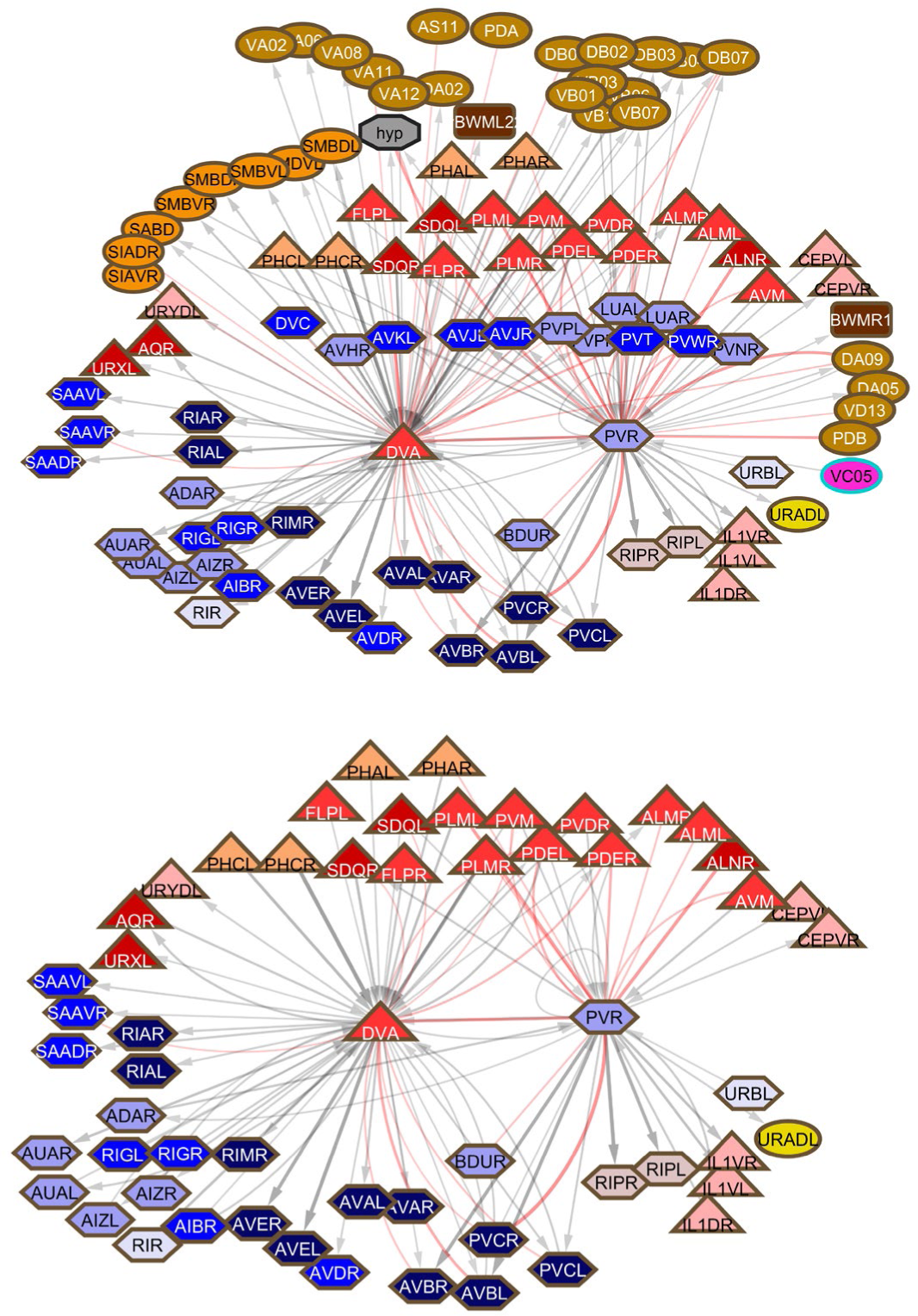
Top: first neighbors of DVA and PVR. Bottom: same with muscles, motor neurons, and ventral cord neurons removed to allow visualization of the large number of sensory inputs.

The touch neuron classes ALM, AVM, PLM and PVM have been long known to target the command interneurons and stimulate a rapid locomotory response (18). However, only a minority of their output connectivity is to these neurons: 11% (33/294 EM serial sections) of the gap junction connectivity and 32% (117/364 EM serial sections) of the chemical output. Along with PVR and DVA, among their additional targets are the BDU neurons, a pair of previously unstudied neurons with lateral cell bodies that send lateral processes into the nerve ring, to which ALM and PLM are strongly connected by gap junctions (19) (**Figure 10**). BDU processes run adjacent to the excretory canal, suggesting a possible sensory function. As is the case for DVA and PVR, both anterior and posterior touch receptors target BDU, indicating this is unlikely to be a signal for locomotory direction. In both sexes, among the nerve ring targets of BDU are sex-specific cells, HSN in the hermaphrodite and MCM in the male. There are also reciprocal connections to the ventral cord interneurons PVN (in the hermaphrodite reconstruction but not the male reconstruction). PVN also has chemical and gap junction connections to HSN, creating a triangular circuit with BDU (see PVN below). In the male, PVN has interactions with many components of the male mating circuitry in the pre-anal ganglion. BDU and PVN are thus apparently involved in circuitry for input from multiple sensory neurons to sexual circuits.

**Figure 10.**
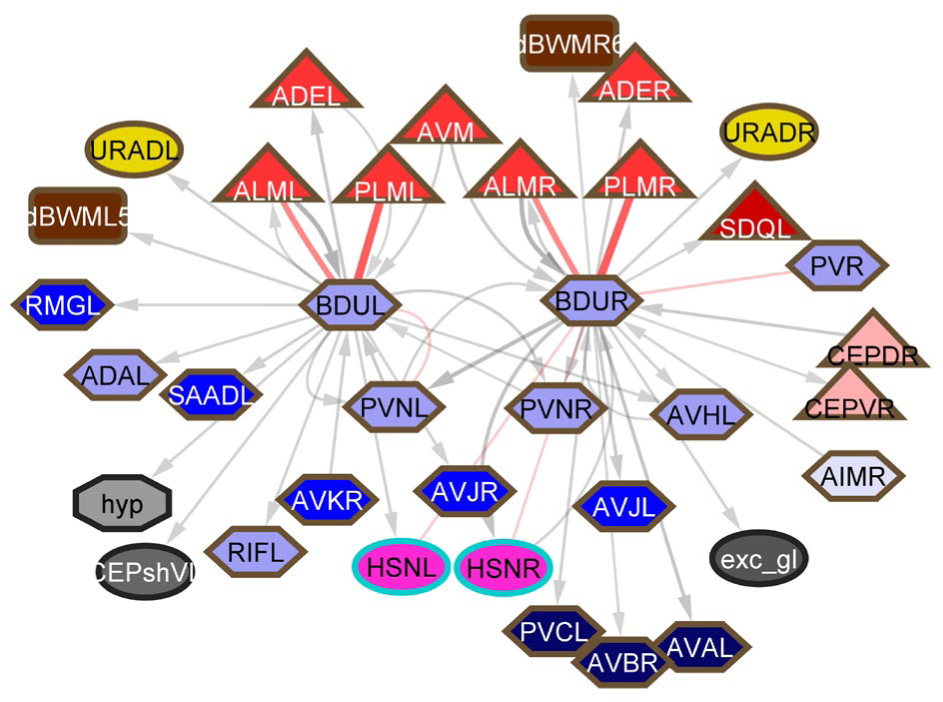
First neighbors of BDU in the hermaphrodite. Bilateral BDU neurons are major targets of the microtubule touch cells.

Two other neuron classes may have an important role in bodywide regulation. Like PVR, ALNL/R and PLNL/R have endings in the tailspike. Rather than sending processes through the ventral cord like PVR, they have processes running to the head in lateral tracts adjacent to the touch cells — ALN in the anterior body region adjacent to ALM, PLN in the posterior body region adjacent to PLM. Like DVA, they have major output onto the sublateral motor neurons (**Figure 11**). They have been reported to have a role in oxygen sensation, but otherwise are unstudied, in spite of their association with the well-studied microtubule touch cells, to which they do not have synaptic connections (except for a gap junction between PLMR and PLNR scored in a single EM section in the hermaphrodite reconstruction) (20). Apart from input from the phasmid neurons, ADE, and PLM, they receive no other sensory inputs and in view of their extensive output onto the sublateral motor neurons, it seems likely that they have an unknown sensory function of their own in addition to oxygen sensation. Notably, in the hermaphrodite reconstruction, there is a significant gap junction connection between ALNR and PVR, thus connecting the DVA/PVR and ALN/PLN networks. This connection is absent in the male reconstruction, which needs to be checked but could be related to the significant reorganization of the adult male tail, which lacks a tailspike.

**Figure 11.**
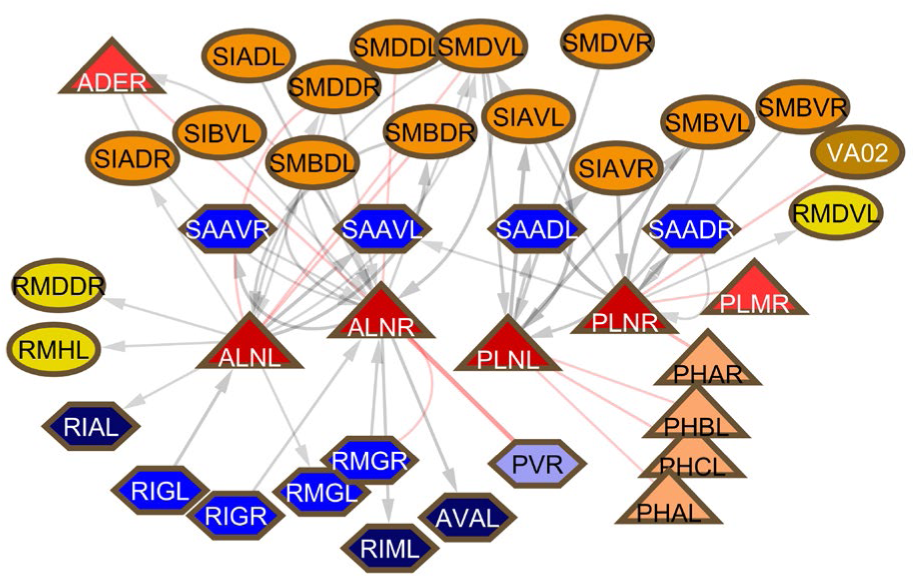
First neighbors of ALN and PLN. SAA neurons have been classified as interneurons and are a major source of input to AVA, but they have lateral processes running into the nose that express genes for stretch receptors and that also make neuromuscular junctions and so in these respects are similar to sublateral motor neurons.

### Functions of the ventral cord neurons

Comparison of their connectivity shows that while they share many connections, each class of non-command interneurons in the ventral cord also has distinct connections (**Supplementary File 8**). Most have some amount of sexually dimorphic connectivity. The differences between them may reflect distinct roles in bringing information from a particular sensory stream into the central network (**Figure 7**), and/or to dispersing information from the central network to a particular point of output. Two aspects of their connectivity, in addition to their module assignment, can be used to gain some information about these separate functions. One is to examine the set of first neighbors in the connectivity diagram. The second is to determine the shortest path between sensory input and motor output on which they lie. However, it should be kept in mind that, as noted, along with distinctive features, each of these neurons also has connections to the other ventral cord neurons including the command interneurons and usually to ventral cord motor neurons and even body wall muscles, emphasizing their importantly distributive nature (**Figure 2B**). It is also important to keep in mind that generalizations drawn from connections or absence of connections, especially weak connections, documented in single reconstructions need to be verified, as they may represent interindividual variation or even reconstruction errors. Missing connections, *e.g.* particularly in the male data in the head, need to be verified.

#### *AVF(L/R)* (comm 10)

AVF and PVQ are two pairs of interneurons involved in sexual circuits. Each has significant sexual dimorphism. Their function in conveying sexual signals to central circuits has been pointed out previously (see Fig 6 in Cook et al. (2)).

##### Hermaphrodite

In the hermaphrodite, in the tail AFV and PVQ are joined by gap junctions. AVF receives chemical input from PHA. In the head, AVF receives chemical input from HSN and from AIM, another interneuron implicated in sexual regulation. Output is to HSN and to AVB. Serotonergic stimulation of AVF by HSN promotes a burst of forward locomotion at the start of a bout of egg laying (21).

##### Male

AVF has male-specific branches in the preanal ganglion and receives significant chemical input from a subset of ray sensory neurons and male-specific interneurons (PVV, PVX, and EF). It is one of three interneuron classes that run through the ventral cord and have output in the head that receive input from the rays: in addition to AVF, these are shared neuron PVN and male-specific interneurons EF1, EF2 and EF3. The spectrum of ray inputs targeting each of these neuron classes is different, suggesting they convey distinctive signaling: most input to AVF (90% — 146/162 EM serial sections) is from the B-type neurons in just four of the rays, those with openings on the dorsal surface of the fan — R1B, R5B, R7B, and R9B. PVN and EF receive input from the B-type neurons in most or all of rays. None receive significant chemical input from the A-type neurons and AVF and PVN have negligible gap junction connectivity, while the EF neurons have a total of 80 sections of gap junctions to three of the A-type neurons — 2A, 6A, and 7A. In the head, AVF has strong, male-specific chemical output to RIF, an interneuron that combines input from head chemosensors via AIA with sexual pathway inputs (2). (The EF neurons but not the PVN neurons also target RIF.) As in the hermaphrodite, AVF has output onto AVB, but in the male the connection is stronger and there is also output onto PVC.

#### *AVG* (comm 9)

##### Hermaphrodite

AVG is not among the high degree neurons (**Fig 6**). Its only significant sensory input is from PHA and PVD. In some animals, it runs all the way through the preanal ganglion and into the tailspike, but no sensory function for it has been documented. It has output onto AVA, AVB, and PVC as well as gap junctions to PHA and DVC. But ablation experiments revealed little to no discernable effect on behavior (22).

##### Male

In the male, in addition to PHA and PVD, AVG receives weak input from a variety of male-specific sensory neurons and interneurons. Most notably, it receives strong chemical input from male-specific sensory neurons HOA and PCA, and shared, sexually dimorphic sensory neuron PHC. Its weak output, both chemical and electrical, is scattered across the same set of neurons from which it receives input. As in the hermaphrodite, this set of output targets suggest no clear role in male behavior (23).

During embryogenesis, AVG pioneers the ventral cord from its cell body in the retrovesicular ganglion (22). It appears that it may provide a similar pioneering or guidepost function postembryonically in the male, where an extensive period of neurogenesis and synaptogenesis from the late L3 to early adulthood establishes the circuits for male mating. All the input from HOA to AVG is at a dyadic synapse with co-recipient PHC. Likewise, the input from PHC to AVG is at dyads with postsynaptic HOA. A similar pattern emerges for HOA connection to PCA: nine of 12 synapses HOA>PCA are at dyads with AVG, while 13/33 synapses PCA>HOA are at dyads with AVG. Unlike the scattered connectivity of AVG, the strong HOA>PHC connection would appear to be important in the mating circuits — HOA senses presence of the vulva and PHC targets important male-specific downstream interneurons PVX, PVY, PVZ and CPn (23) (NB, in Jarrell et al. (23), PHC is misidentified as LUA). The reciprocal connections between HOA and PCA join two male-specific sensory neurons that detect the vulva. The many synapses between HOA and sex shared neuron PHC occur along a male-specific process extended by PHC along AVG during the L4 larval stage (24). Ablation of AVG attenuates this outgrowth. HOA extends a process that is required to find this growing PHC process. Strikingly, although they eventually run together extensively, HOA and PHC develop their reciprocal presynaptic densities only when AVG is also present as co-recipient, as if AVG is necessary for formation of the HOA<>PHC connection. AVG may thus serve as a landmark internal to the preanal ganglion for assembly of parts of the male mating circuits.

Singhvi and Shaham (25) have pointed out the many similarities between *C. elegans* glial cells and astrocytes. AVG, viewed heretofore as a neuron, shares some of these glial cell properties. It expresses UNC-6/Netrin to facilitate its guidance function for the ventral cord, while its presence at HOA and PHC synapses resembles the tripartite astrocyte synapse and similar structures made by the *C. elegans* CEPsh glial cells (26). In several places in the male preanal ganglion, AVG is striking and unique in extending processes that surround other neurons (unpublished observations). AVG has some synapses, is cholinergic, and does not express glial-specific genes (S. Shaham, personal communication). It is therefore perhaps best viewed as a hybrid cell type with both neuronal and glial-like properties.

#### AVH(L/R) (comm9)

##### Hermaphrodite

AVH, which is connected to both AVF and PVQ by gap junctions and reciprocal chemical connections, appears to be somewhat similar to them in overall connectivity. For example, like PVQ, AVH is connected by gap junctions to chemosensory neurons ASK and PHB. It is also connected by gap junctions to two posterior motor neurons, AS11 and VD12, as is AVF. It receives weak chemical inputs from sensory neurons in the head. What distinguishes AVH from the other ventral cord neurons, including AVF and PVQ, is chemical output to RIR (comm 3) and sublateral motor neurons SMB. As described below, RIR aggregates information from a variety of sensory neurons and targets important interneurons AIZ in comm 3 and RIA in comm 4, creating many triangular circuits. Thus one role for AVH could to be to contribute input to this information stream from, for example, ASK, which is otherwise not connected to AIZ or RIA: ASK>AVH>RIR>AIZ,RIA. (PVQ does not target RIR, AIZ or RIA.)

##### Male

The gap junctions to ASK, PHB, AS11 and VD12 are absent and there is a strong electrical connection to PHA. PVQ also has a strong, male-specific electrical connection to PHA. Otherwise, AVH connections are the same as in the hermaphrodite, but weaker; for example, there is only weak connectivity to RIR. There is scattered and weak input from several male-specific neurons in the tail.

#### *AVJ(L/R)* (comm 9)

##### Hermaphrodite

The little chemical sensory input is from ADL, AQR, PQR, FLP and URX (all 0_2_/aversive inputs?). Distinctive chemical input is from PVR (comm 9) (discussed above and see below). An additional distinguishing feature is five gap junctions to RIS (comm 4). GABAergic RIS appears to distribute a presumptively inhibitory signal to sensory, inter, and motor neurons of comm 4 (see below).

##### Male

Input from ADL and ADA are present as in the hermaphrodite, but otherwise most sensory and interneuron inputs, including those from PVR, are absent. Likewise, the large number, albeit weak, of gap junction connections present in the hermaphrodite are also absent, including the connection to RIS. Thus AVJ may be synaptically less active in the male, but the possibility of incomplete male reconstruction should be kept in mind.

#### *AVK(L/R)* (comm 9)

##### Hermaphrodite

AVK is the primary target of sensory neuron PDE (receiving 50% of PDE output by weight) and also receives input from AVM and PVM. It is distinctive in receiving chemical input from RIS (comm 4), RIG (comm 4), and RMF (comm 9). There is possible sensory input via gap junctions from DVA and AQR. Distinctive output is weak chemical connectivity to three neuron classes of the head motor system, SAA and RIM in comm 3 and RIV in comm 4, and to all of the sublateral motor neurons except SAB. There is unique electrical connectivity to SMB. RIV, SAA, and SMB are part of a turn circuit that inhibits reversals (27). Thus, it would seem one role of AVK is to aggregate several diverse streams, both sensory and interneuronal, and connect these to sublateral motor neurons and this turning circuit. AVK receives chemical input from and is connected via gap junctions to the unstudied high-degree hub-and-spoke neuron RIC (see below).

##### Male

Connections are the same as in the hermaphrodite, except that there is a strong electrical connection to PVP, over some 53 serial sections, whereas in the hermaphrodite there is a gap junction in just a single section. The function of this sexually dimorphic connectivity is unknown. Chemical outputs and the remaining gap junction connections are to the same set of neurons as in the hermaphrodite, but even weaker.

#### *AVL* (comm 9)

##### Hermaphrodite

AVL functions in defecation, where it is partially redundant with DVB, which also runs part way in the ventral cord, in controlling the defecation cycle (28). It has stimulatory GABAergic output onto the intestine and gap junctions to several D-type (inhibitory) ventral cord motor neurons in the posterior. Extensive input and output connectivity across the nervous system and nerve ring attests to the integration of defecation behavior with other behaviors.

##### Male

In the male, the functions of both AVL and DVB are diverted to the copulatory circuits, consistent with the fact that the anal opening is now a cloaca that must also accommodate the expulsion of gametes (see Fig 6 of Cook et al (2))(29). Chemical synapses onto the intestine are not present. The gap junctions to the D-type inhibitory motor neurons are absent and instead there are 30 sections of gap junctions onto PDB, a likely excitatory cholinergic AS-type motor neuron that has neuromuscular junctions to dorsal body wall muscles in the posterior.

#### *DVA* (comm 9)

##### Hermaphrodite

DVA, like PVR, to which it is connected by gap junctions, has chemical inputs from the family of proprioceptive neurons of all types across the body and itself has a mechanosensory stretch response (1) (**Figure 9**). It has chemical output across a spectrum of interneurons, command interneurons, sublateral and ventral cord motor neurons. Its apparently important role in the nervous system is reflected by high degree in both synaptic and peptidergic networks as discussed above.

##### Male

Chemical inputs are from the same set of proprioceptive neurons with the exception that input from PHC is absent, possibly reflecting the diversion of PHC into the copulatory circuits. Otherwise, circuitry is the same as in the hermaphrodite, with the possible exception that chemical output onto ring interneuron RIR is far stronger in the male reconstruction.

#### *DVC* (comm 9) and *PVT* (comm 9)

##### Hermaphrodite

DVC and PVT, connected by gap junctions, have such similar connectivity that they may be considered in this respect to be a neuron pair, even though they have unrelated lineal origins: both are embryonic, but DVC is from the C blastomere while PVT is from ABp (see **Supplementary File 8** to compare the connectivity). The processes that each sends anteriorly from its posterior cell body (DVC in the retrovesicular ganglion, PVT in the pre-anal ganglion) run together through the ventral cord and remain in contact as both progress around the nerve ring. Neither has significant sensory input but a sensory function for DVC has been documented — a stretch receptor function that stimulates backwards locomotion through chemical connections to AVA (30). They share chemical output to several navigational interneurons, including, notably, RIG, with a single exception: DVC targets AVA but PVT does not. Both neurons are so highly connected to other ventral cord neurons by gap junctions, particularly PVP, that their influence must be considered widespread. PVT, as noted above, is a hub of the neuropeptide communication network (5).

##### Male

The connections in the hermaphrodite are present in the male with, again, the single exception that DVC does but PVT does not target AVA, so this difference is unlikely to be a reconstruction artifact. DVC has scattered, weak chemical input from and electrical connections to a number of male-specific neurons in the tail circuits that are not shared by PVT, while PVT has some input from male-specific sensory neuron CEM in the head not shared with DVC. Both neurons make gap junctions to male-specific sensory neuron SPV, which is involved in ejaculation.

#### PVN(L/R) (comm 9)

##### Hermaphrodite

PVN is a high-degree neuron like the other ventral cord neurons, but its interactions are so diverse (for example, interactions with ventral cord motor neurons and body wall muscles mostly in the head but some also in the tail) and so weak that it is difficult to discern a specific role. The exceptions are unique reciprocal chemical and electrical connections to BDU. As discussed above, BDU receives chemical input from ALM, but an unknown function, possibly sensory or physiological, is suggested by the presence of a process extending down the body next to the excretory canal. Chemical input from BDU is greater than output to BDU, suggesting one role of PVN may be to convey the BDU signal to the central network. Both BDU and PLN have interactions with the sexual neurons HSN and VCn.

##### Male

Connections to BDU are absent. As in the hermaphrodite, there are interactions with sex-specific circuitry. There is significant chemical input from the rays: 164 sections almost exclusively from the B-type neurons in every ray except ray6. Output is to the same ray B-type neurons and to the male-specific interneurons EF, PVV, PVX, and PVY, and to AVB (including one 16 section gap junction between PVNL and AVBL). Thus PVN is somewhat like AVF in collecting input from the rays and directing output to EFn and AVB, but as noted above, the subset of input ray neurons is different and whereas AVF connections to AVB are mostly in the head and to EFn in both head and tail, the PVN synapses to EFn and AVB are all in the tail.

#### *PVP(L/R)* (comm 9)

##### Hermaphrodite

PVP has the highest gap junction degree of any neuron in the nervous system. Among these gap junction connections, the most notable are connections to the pair of sensory neurons with sensory endings facing the coelomic cavity, AQR in the head (102 sections) and PQR in the tail (26 sections), and to the neuron pair DVC (54 sections) and PVT (31 sections). There is little input via chemical synapses. The main chemical output, in addition to connections to other central network neurons, is to AVA, AVB, and PVC. AQR and PQR also target AVA, AVB, and PVC, thus creating a triangular circuit including PVP. There is some presynaptic chemical connectivity of PVP to RIG(L/R) (comm 4). DVC and PVT also target RIG, creating another triangular circuit with PVP. This RIG connectivity is notable because RIG aggregates input from several sensory neurons, including oxygen sensors and URX. URX, like AQR and PQR, has sensory endings facing the coelomic cavity, but RIG has no input directly from AQR or PQR. Conveying additional sensory input to RIG may be a role of PVP. PVP is involved in regulating the pattern of locomotion, roaming versus dwelling (31). It appears to develop hermaphrodite-specific branches that have wing-like sensory endings surrounding the egg-laying apparatus at the vulva. PVP might thus play a role in regulating egg-laying or locomotion during egg-laying (32).

##### Male

There is no clear sexual dimorphism of the connectivity. The gap junctions to AQR, PQR, DVC and PVT are present but not as strong as in the hermaphrodite. Likewise there is chemical output to AVA and AVB (but not PVC) and to RIG, but all weaker than in the hermaphrodite. There are no apparently significant interactions with the male-specific tail circuits.

#### *PVQ(L/R)* (comm 10)

##### Hermaphrodite

The relatedness to AVF is noted above (and see circuit diagrams in Fig 6 of Cook et al. (2)). PVQ is joined to AVF by gap junctions in the tail and like AVF receives input from PHA. PVQ also receives input from PHC and there is a weak electrical connection to PHB. A distinctive feature of PVQ is left right homologs are strongly connected to each other in the preanal ganglion by gap junctions. In the head PVQ is connected to two pheromone sensors: chemical input from ASJ and gap junctions to both ASJ and ASK. In addition to reciprocal chemical output to ASJ and ASK, the main output is to AIA, an interneuron targeted by many amphid sensory neurons, including ASK but not ASJ. Thus there is a feedforward loop incorporating PVQ connecting ASK to AIA, but connectivity from ASJ to AIA is solely via PVQ.

##### Male

In the male head, as in the hermaphrodite, there is chemical input from ASJ and electrical connectivity to ASK and chemical output to AIA as well as to AVF. In the tail there is chemical input from the EF class of male-specific interneurons and some weak input from PHA and PHB. There is electrical connectivity to male-specific interneurons CA05 and CA06. In a major sexual dimorphism, there is a strong gap junction connection to PHA (70 EM sections) that is absent in the hermaphrodite. This creates a one-neuron electrical connection between PHA in the tail and pheromone sensor ASK in the head.

#### *PVR* (comm 9)

##### Hermaphrodite

The apparent role of PVR, a possible mechanosensory neuron with extension into the tail whip, as a hub of a bodywide sensory network, its connection to DVA and together with DVA its status as a hub neuron of the neuropeptide connectome, and its output onto the pharyngeal regulatory interneuron RIP, is described above (**Figure 9**). These properties appear to lend to PVR a significance in the overall function of the nervous system that has been previously unrecognized.

##### Male

Absence in the male reconstruction of a gap junction connection to ALNR, which links the DVA/PVR network to the ALN/PLN network, needs to be confirmed, but could be related to the fact that there is no tailspike in the male. Otherwise, connectivity is the same as in the hermaphrodite, so this system is not sexually dimorphic.

#### *PVT* (comm9)

See DVC.

### Functions of ring interneurons

A number of neurons have been classified as ring interneurons because their processes are contained entirely within the nerve ring (3). Some of these have properties similar to the ventral cord neurons discussed above — they have high degrees, a large number of gap junction connections, and are understudied. Like the ventral cord neurons, several target muscles and so have been classified previously as motor neurons. As noted above, several are distinguishing targets of the non-command ventral cord interneurons. Below are deductions regarding functions of a subset of these neurons.

#### RIC(L/R) (comm 4)

RIC, an octopaminergic neuron, is one of two ring interneurons, along with RIR, that receives inputs from several sensory neurons and targets many of the same neurons as those sensory neurons, creating triangular circuits (**Figure 12**). Octopamine is expressed by RIC in the absence of food (33). Dopamine signaling from one of the connected sensory neurons, CEP, suppresses expression in the presence of food. This suggests that the regulatory role of RIC in the triangular circuits is related to the response to food. RIC has no significant interaction with non-command ventral cord interneurons.

**Figure 12.**
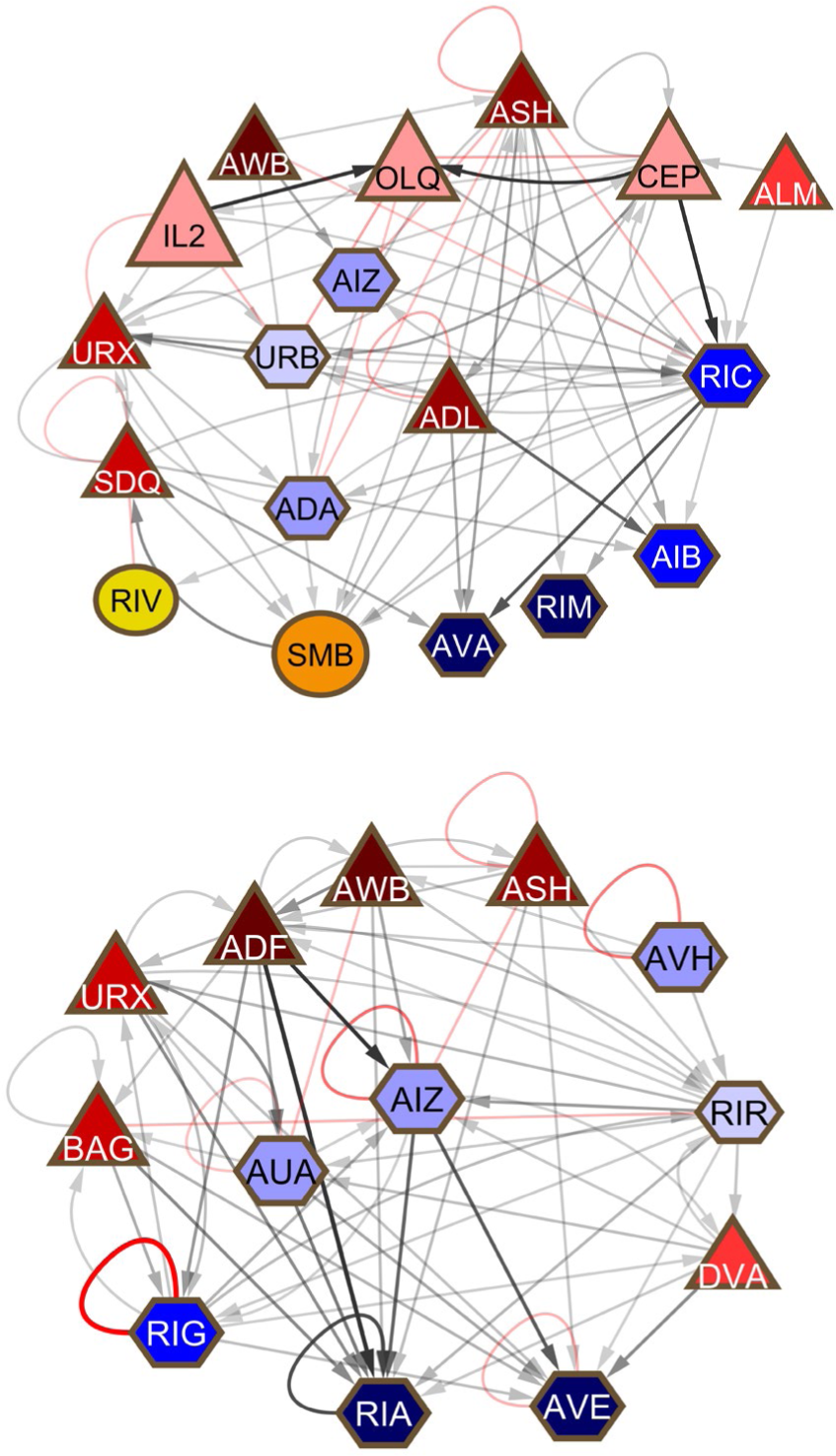
Two ring neurons, RIC (*upper*) and RIR *(lower*) lying on multiple triangular pathways between sensory neurons and their targets.

#### *RIF(L/R)* (comm 9)

The connectivity of RIF implicates it as a nexus of sexual signals and somatic signals relevant respectively to reproduction and behavior. This has been pointed out previously (2). Somatic signaling comes through AIA. Sexual signaling is from AVF in both sexes, HSN in the hermaphrodite, and, in the male, two classes of male-specific interneurons, MCM and EF. RIF expresses receptors for two sex-promoting signals, PDF and nematocin, and lies on a functional pathway between sex pheromone and reproductive behavior and physiology (34–36).

#### *RIG(L/R)* (comm 4)

RIG and RMG are two high-degree neurons in comm 4 that have an overall similar pattern of connections. They receive inputs from a large number of sensory neurons, often by gap junctions, and have output onto a spectrum of downstream targets (**Figure 13**). Unlike RMG (see below), RIG has not been studied and has no documented function

**Figure 13.**
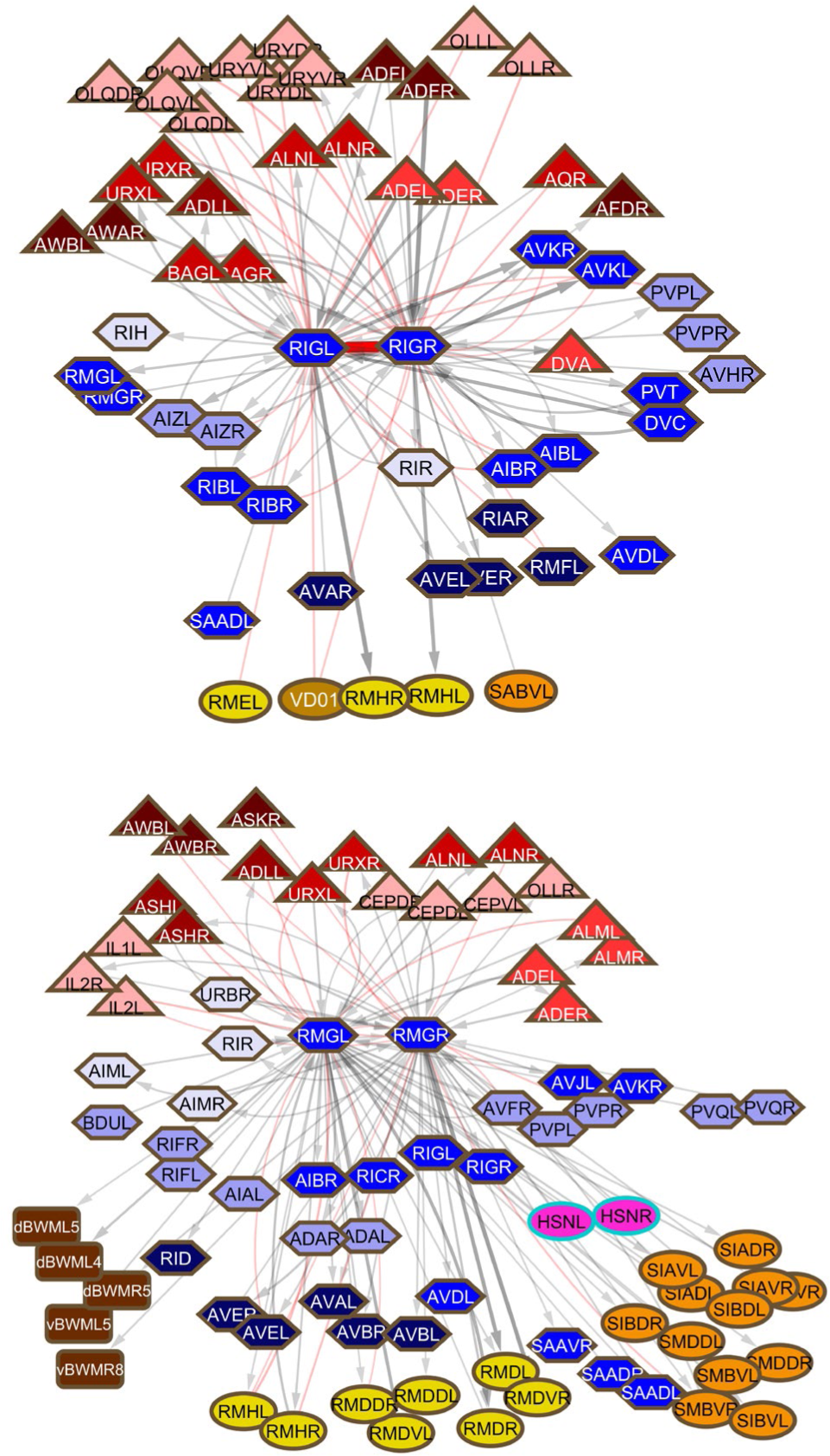
Hub-and-spoke neurons RIG (*upper*) and RMG *(lower*). The pattern of a large number of gap junction connections to sensory neurons and a spectrum of output targets is similar for these two high degree neurons, but the sets of cells involved are largely non-overlapping.

presently, but the similarity to RMG suggests this may be considered a hub- and-spoke neuron. RIG and RMG are connected to each other and have multiple connections to non-command ventral cord interneurons, implicating them in a widespread role. Notably, RIG makes both chemical connections to and has gap junction connections with AVK.

#### *RIR* (comm 3)

RIR, an unstudied ring interneuron, is placed at the top of the hierarchical network in layer 4 by the layering algorithm (2). Like the other neurons in this group, it makes very few gap junctions, the exception being gap junctions to oxygen-sensing BAG neurons. RIR connectivity resembles that of RIC (see above) in creating triangular pathways involving sensory neurons and their targets (**Figure 12**). Its presumptive regulatory role in these circuits is unknown. In interactions with the central network of ventral cord non-command interneurons, RIR receives input from AVH and has reciprocal interactions with DVA and PVP.

#### *RIS* (comm 4)

RIS is GABAergic and hence presumptively inhibitory. It has only scattered, weak chemical input from sensory neurons and interneurons across the network. Sensory input is from proprioceptors SDQ and FLP, but there is greater output than input to CEP, URY, and OLL. Chemical output and gap junctions are to the head motor system, RIM, AVE, RMD, and SMD. The most prominent connection is gap junction connectivity over 15 EM sections to AVJ, which also makes a weak chemical connection to RIS. This AVJ connectivity would appear to bring input from comm 9 to an inhibitory signal within comm 4 — CEP, URY, OLL, AVE, RMD, and SMD are all in comm 4. RIS is required for developmentally timed sleep (37).

#### RMG (comm 4)

RMG, the neuron with the highest chemical outdegree, has been studied in some detail (38). RMG is the hub of a “hub-and-spoke” circuit that aggregates via gap junctions information from several sensors involved in regulating social behaviors (worm aggregation, responses to oxygen and pheromones) and has output onto elements of the motor system, particularly in the head (**Figure 13**). RMG activity is regulated by activity of the neuropeptide Y receptor homolog gene *npr-1*, revealing how a set of connections is coordinately regulated by a neuropeptide. RMG is connected to RIC (see above) and like RIC connects at multiple points to the network of non-command ventral cord interneurons. It is classified in White et al. (3) as a motor neuron due to its muscle connectivity.

## Discussion

Although an anatomical description of the synaptic connectivity of the *C. elegans* nervous system has been available for nearly forty years, providing the basis for many genetic and experimental investigations, the functions of many of the neurons remain poorly documented or are unknown (3). A recently available updating and completing of the connectome across the entire body, including all end organ connectivity, provides an opportunity to query the functions of neurons (2). This approach was used previously to assign functions to the male-specific set of neurons in the male tail that govern mating behavior (23).

Not only does the connectome provide insight into the functions of individual neurons, it also makes possible a perspective on overall nervous system architecture. One characteristic that emerges is the significant role played by the end organs themselves. End organs are simultaneously at the bottom and at the top of the hierarchical structure — at the bottom they are important points of convergence for the many pathways from sensory inputs, and at the top, their output affects those inputs, creating a feedback loop. While the body wall musculature is considered here, there are without doubt major effects of the output of other physiological and reproductive system functions as well. First, the body wall muscles are connected by gap junctions so that their activity is affected by the activity of their neighbors. Second, individual muscle cells are points of circuit convergence (**Figure 2**). Third, connectivity has uncovered a bodywide system of mechanosensors converging on two singleton interneurons, DVA and PVR, which then disperse their outputs widely (**Figure 9**). In addition to these two neurons, mechanosensors have additional targets beyond the well-studied command interneurons that activate locomotion. The body surface thus emerges as a large and important receptive field with targets throughout the nervous system. The output of this sensory field will be directly affected by the contractions of the body wall muscles.

A second characteristic that emerges is the apparent importance of gap junction connections, especially for certain classes of neurons (**Figure 6**). In the adult hermaphrodite reconstruction of the non-pharyngeal nervous system, 21% of the connections by number (number of edges in the graph) (1241/5808) and 19% of the connectivity by weight (total weight of edges in EM serial sections) (6481/32421) are gap junctions (2). (The very large amount of scored gap junction connectivity involving CAN, exc_cell, hmc, and hyp is excluded from this calculation.) The disproportionately large number of gap junctions scored among the posterior circuits (**Figure 8**), reported here and not noted previously, raises the possibility that this is an artifact of the fixation, imaging, or scoring: either under-scoring in the anterior series or over-scoring in the posterior ones. This possibility is mitigated by the fact that the anterior series in the hermaphrodite, N2U, and the posterior series in the hermaphrodite, JSE, and male, N2Y, were prepared and imaged by the same individual during the same period (J. N. Thomson, MRC Cambridge laboratory). On side-by-side comparison, the electron micrographs look similar. Evidence for the validity of the scored gap junctions comes from the consistency of these connections in the connectome. For example, left/right homologs frequently create gap junctions to the same target or left/right homologous targets. Many other, sometimes striking, examples may be noted. For instance, PVP makes consistent, strong, gap junction connections in the nerve ring to AQR and in the tail to the neuron considered to be the AQR equivalent, PQR, in both sexes. This result involves four separate animals and EM series scored by two different individuals. Within the N2Y series, PVQL is joined to PHAL and PVQR is joined to PHAR, but neither is joined as heavily to the many other processes that they contact. The distribution among neurons is uneven, even though their amount of neighbor contacts is similar (**Figure 6**). Where a contactome is available from volumetric tracings, there is no correlation between the amount of contact and the number of gap junctions, as might be expected for a fixation or staining artifact (this laboratory, unpublished). Finally, in behaving animals, the activities of neuron pairs connected by gap junctions in the EM-based connectome is more highly correlated than the activities of pairs connected by chemical synapses (39, 40). These observations suggest that, despite the uncertainty often felt in confidently identifying them in electron micrographs, scored gap junctions in *C. elegans* connectomes are reliable. If this is true and the relatively lesser number scored in the anterior series is a reconstruction artifact, the artifact would be under-scoring in the N2U series. In this case, the nervous system has a still higher level of gap junction connectivity than currently realized.

An important feature of overall nervous system architecture is the way multiple information streams are brought together for computing output — multisensory integration. The number of sensory inputs far exceeds the number of possible outputs. Strikingly, in *C. elegans*, more than half of the neurons in the somatic nervous system of the hermaphrodite (53% percent, 149/280) have demonstrated or possible sensory function. Included in this number are several neurons classified as interneurons or motor neurons, but which also have a sensory function. The most important of these are the ventral cord motor neurons, which have stretch-sensitive dendritic extensions (41). Three classes (twelve neurons) that run laterally making neuromuscular junctions, SAA, SMB, and SMD, express genes for known stretch receptors (42). Also are included DVC, PVR, BDU(L/R), and PVP(L/R). Two members of the “UR” set, URA (motor neuron) and URB (interneuron) are also included: “UR” stands for “unknown receptor” because these neurons have apparent dendritic extensions towards the nose similar to many other sensory neurons, including URX and URY (J. White, personal communication). Considering computation for locomotion and posture, in a massive process of convergence, information originating from these sensors is aggregated to specify a single scalar quantity in each muscle cell, the muscle tension generated.

The connectome reveals that multisensory integration occurs throughout the network. The dispersed nature of the information processing is reflected in most interneurons having outdegrees equal to, and in some cases even greater than, their indegrees. Perhaps surprisingly, many information streams are brought together at the very last step, where each muscle cell combines inputs from an average of ten neurons. For just one modality, chemosensation, some 7% of the genes in the genome are putative chemoreceptors of the seven-transmembrane G-protein-coupled receptor class (1280 genes) (43). Apparently, the concentration of each of over 1000 compounds is evaluated and compared to the concentrations of all the others as input relevant to decision-making. The far larger number of chemoreceptor proteins than chemosensory neurons, as well as the polymodal capacities of some neurons (for example the nociceptive ASH neuron is polymodal for osmo-, mechano-, electro-, photo- and odorsensation) means much of the integration of incoming sensory information inevitably occurs within the sensory neurons themselves. Immediately downstream, circuit mechanisms have been studied that involve connections between sensory neurons, connections of sensory neurons to dedicated interneurons (such as the amphid interneurons AIA, AIB, AIZ and AIY, and the hub interneuron RMG), and showing how these interactions may be affected by neuromodulators (44). But the connectivity reveals convergence occurs throughout the network right down to single muscle cells.

Among the new findings, a previously unrecognized locus of information processing appears to be a network of high degree neurons running in the ventral cord and connected widely throughout the nervous system (**Figure 7**). Remarkably, these neurons are among the most heavily electrically coupled and some are hubs in the extra-synaptic, peptidergic communication network. They are among the least studied neurons in the nervous system. The nerve ring neuropil has always been considered the nematode “brain.” John White has pointed out it closely resembles a somatotopic brain region, where sensory/motor connections are arrayed physically in congruence with motor output (45). The balance of information processing between that which occurs in the nerve ring and that which occurs outside it within the central network of ventral cord neurons and elsewhere remains to be seen (**Figure 8**). The large amount of sexual dimorphism in the ventral cord group may reflect a central role.

Along with the function of the ventral cord neurons, previously unrecognized significant functions of several additional neurons are revealed by connectivity. These include the eight neurons with processes extending into the tailspike of the hermaphrodite, ALNL/R, PLNL/R, PHCL/R, PVR and AVG. While the function of the tailspike or whip has never been studied and a proprioceptive function for these neurons remains speculative, connectivity suggests important circuit functions for five of them. As mentioned above, PVR, together with another ventral cord neuron DVA, to which it is connected, appears to function as an integrating interneuron of a sensory system whose receptive field is the body surface (**Figure 9**). ALN and PLN target the sublateral motor neurons and contribute 20% of the chemical input to SAA, a class of four neurons also with lateral processes making neuromuscular junctions similar to the sublateral motor neurons (but unlike the other sublaterals, has significant chemical output onto AVA) (**Figure 11**). ALN and PLN have been implicated in oxygen sensation and receive sensory input from phasmid neurons, but it seems likely they have additional sensitivities. In the hermaphrodite reconstruction, there is a significant gap junction connection between PVR and ALNR.

An unexpected finding was the near identity of connectivity of DVC and PVT. Curiously, this seeming oddity of pairing a cell descended from embryonic blastomere AB.p (PVT) with one descended from the C blastomere (DVC) is shared with the pair DVA PVR — DVA is descended from AB.p while PVR is descended from C. PVR and DVC are lineal first cousins and the only neurons produced by the C lineage (which otherwise generates hypodermal and muscle cells). PVT shares properties with another singleton, AVG, in expressing UNC-6/netrin and having a guidepost role in development and maintenance of the ventral cord (26, 46). The finding of similar or related synaptic connections of pairs of neurons, like PVR and DVA, and DVC and PVT, suggests investigation of the phenotypes of the double ablations.

While the extensive connectivity of the ventral cord neurons indicates they may influence many neural pathways, their unique or distinctive connections suggest circuit-specific roles. Noteworthy among these are the robust gap junction connections of PVP to the pseudocoelom sensors AQR and PQR and the ventral neuron pair DVC and PVT. PVP, DVC and PVT target RIG, which receives direct input from the other pseudocoelom sensor URX. This might be a pathway aggregating multiple sensory inputs from the body, including from possible additional sensory modalities of DVC and PVT. What this would have to do with a function of PVP in regulating locomotion during egg laying is unclear and illustrates the potentially widespread roles of extensively and electrically coupled neurons such as PVP. Additional examples of suggested pathways and specific interneuron functions are the connections of ASJ and ASK to PVQ, ADL to AVJ, PDE to AVK, and BDU, a gap junction target of touch neurons, to PVN. All these relationships and many others indicated by the connectivity suggest directions for future research.

## Materials and Methods

The analysis is based on the data of Cook et al., (2). The cytoscape files that are the basis of the figures in that paper are available at WormWiring.org. Connectivity diagrams were prepared from these files employing the network analytical features of Cytoscape. Indegree and outdegree values were determined from the Excel file adjacency matrices of Cook et al. (2) Graph analysis for community detection and betweenness centrality was carried out with a MATLAB package prepared by Adam Bloniarz and available at WormWiring.org.

## Supporting Information

Supplementary files are submitted separately.

## Acknowledgements

I am grateful for their comments on the manuscript to H. Buelow, D. Hall, O. Hobert, P. Kurshan, and J. White. This work was supported by NIH grants from NIHD (P30HD071593 to S.W.E.), NIMH (R01MH112689 to S.W.E.), and NIGMS (R01GM066897 to S.W.E.).

