## Supplementary File 3 for "Functions of *C. elegans* neurons from synaptic connectivity"

chem = AdjacencyMatrix('file', 'Herm chem.csv')

chem =

AdjacencyMatrix object of type unknown

300 presynaptic cells

444 postsynaptic cells

4879 entries in adjacency matrix

>> getCommunitiesMM(chem)

Modularity of community structure: 0.515

Community 1

DA02, DA03, DB02, AS03, VD03, dBWML10, dBWMR12,

Community 2

URXR, PLNR, ADEL, ADER, IL2L, IL2R, CEPDL, CEPDR, CEPVL, CEPVR, URYDL, URYVL, URYVR, OLLL, OLLR, OLQDL, OLQDR, OLQVL, OLQVR, IL1R, RIH, URBR, AUAR, RMGL, RMGR, RICL, RICR, RIAL, RIAR, AVER, RMDDL, RMDDR, RMDL, RMDR, RMDVL, RMDVR, RIVL, RIVR, RMHL, RMHR, SMDDR, SMDVL, SMDVR, SMBVR, SIBDR, SIAVL, SIAVR, pm1, pm2D, pm2VL, pm3D, pm3VL, pm6D, pm6VL, pm6VR, pm7D, pm7VR, mc3V, e3D, e3VL, e3VR, g2L, g2R, dBWMR2, vBWML5, vBWML7, vBWMR5, vBWMR6, vBWMR7, vBWMR24, GLRVL, mu_intL, mu_intR, vm1AR,

Community 3

PLNL, IL2DL, IL2DR, IL2VL, IL2VR, IL1DL, IL1DR, IL1L, IL1VL, IL1VR, RIPL, RIPR, URADL, URADR, URAVL, URAVR, RMEL, RMER, RMED, RMEV, SABD, SABVL, SABVR, SMBVL, pm3VR, dBWML3, dBWML4, dBWML6, dBWMR1, dBWMR3, dBWMR4, dBWMR6, vBWML1, vBWML2, vBWML3, vBWML4, vBWML6, vBWMR1, vBWMR2, vBWMR3, vBWMR4, exc_gl, hyp,

Community 4

ALNR, SDQR, URYDR, SAAVR, SMDDL, SMBDL, SIBDL, SIADL, dBWMR5, dBWMR7, dBWMR9, dBWMR10, dBWMR11,

Community 5

SDQL, SMBDR, SIADR, dBWML2, dBWML5, dBWML9, dBWML12, dBWML13,

Community 6

M5,

Community 7

MCL, MCR, mc2DL, mc2DR, mc2V,

Community 8

CEPshDL, CEPshVL,

Community 9

ASIL, ASIR, AWAL, AWAR, AWBL, AWBR, ASEL, ASER, ADFL, ADFR, AFDL, AFDR, AWCL, AWCR, BAGL, BAGR, URXL, ALNL, AINL, AINR, URBL, RIR, AIYL, AIYR, AUAL, AIZL, AIZR, RIS, ADAL, ADAR, RIBL, RIBR, RIGL, RIGR, AIBL, AIBR, SAADL, SAADR, SAAVL, AVKL, AVKR, DVC, PVT, RIML, RIMR, AVEL, RMFL, RMFR, VA01, VB01, CEPshVR,

Community 10

ASJL, ASGL, ASKL, AIML, AIAL, PVQL,

Community 11

AVL, SIBVL, SIBVR, DA01, DA04, DB01, DB03, AS01, AS02, AS04, AS05, DD01, DD02, DD03, VA02, VA03, VA04, VA05, VA06, VA07, VB02, VB03, VB04, VB05, VD01, VD02, VD04, VD05, VD06, VC01, VC02, VC03, dBWML7, dBWML8, dBWML11, dBWMR8, vBWML8, vBWML9, vBWML10, vBWML11, vBWML12, vBWML13, vBWML14, vBWML15, vBWMR8, vBWMR9, vBWMR10, vBWMR11, vBWMR12, vBWMR13, vBWMR14, vBWMR15,

Community 12

DVB, DA05, DA06, DA07, DA08, DA09, PDA, DB04, DB05, DB06, DB07, AS06, AS07, AS08, AS09, AS10, AS11, PDB, DD04, DD05, DD06, VA08, VA09, VA10, VA11, VA12, VB06, VB07, VB08, VB09, VB10, VB11, VD07, VD08, VD09, VD10, VD11, VD12, VD13, VC06, dBWML14, dBWML15, dBWML16, dBWML17, dBWML18, dBWML20, dBWML21, dBWML22, dBWML23, dBWML24, dBWMR13, dBWMR14, dBWMR15, dBWMR16, dBWMR17, dBWMR18, dBWMR19, dBWMR20, dBWMR21, dBWMR23, dBWMR24, vBWML16, vBWML17, vBWML18, vBWML19, vBWML20, vBWML21, vBWML22, vBWML23, vBWMR16, vBWMR17, vBWMR18, vBWMR20, vBWMR21, vBWMR22, vBWMR23, int,

Community 13

I1L, I1R, I2L, I2R, I3, I4, I5, I6, M1, M2L, M2R, M3L, M3R, M4, MI, NSML, NSMR, pm4D, pm4VL, pm4VR, pm5D, pm5VL, pm5VR, g1p, g1AL, g1AR, bm,

Community 14

dBWML19, vBWMR19,

Community 15

ASJR, ASGR, ASKR, ASHL, ASHR, ADLL, ADLR, AQR, PQR, ALML, ALMR, AVM, PVM, PLML, PLMR, FLPL, FLPR, DVA, PVDL, PVDR, PDEL, PDER, PHAL, PHAR, PHBL, PHBR, PHCL, PHCR, AIMR, AIAR, ALA, PVQR, RIFL, RIFR, BDUL, BDUR, PVR, AVFL, AVFR, AVHL, AVHR, PVPL, PVPR, LUAL, LUAR, PVNL, PVNR, AVG, AVJL, AVJR, AVDL, AVDR, PVWL, PVWR, RID, AVBL, AVBR, AVAL, AVAR, PVCL, PVCR, HSNL, HSNR, VC04, VC05, CANL, CANR, exc_cell, vm2AL, vm2AR, vm2PL, vm2PR,

Community 16

dBWML1, dBWMR22, CEPshDR, GLRDR, GLRR, GLRVR,
