## Supplementary File 4 for "Functions of *C. elegans* neurons from synaptic connectivity"

>> gap = AdjacencyMatrix('file', 'Herm gap.csv')

gap =

AdjacencyMatrix object of type unknown

462 presynaptic cells

462 postsynaptic cells

2891 entries in adjacency matrix

>> getCommunitiesMM(gap)

Modularity of community structure: 0.646

Community 1

ASJL, ASJR, ASKL, ASKR, ASHL, ALNL, PLNL, PLNR, AQR, PHAL, PHAR, PHBL, PHBR, PHCL, AIMR, PVQL, PVQR, AVFL, AVFR, AVHL, AVHR, PVPL, PVPR, AVG, DVB, RICL, SAADL, AVKL, AVKR, DVC, PVT, AVL, SMBDL, SMBDR, SIADL, AS11, DD05, DD06, VD10, VD11, VD12, VD13, VC05, VC06,

Community 2

PVDL, PVDR, ALA, AS03, CANL, CANR, exc_cell, hyp,

Community 3

dBWML13, dBWML14, dBWML15,

Community 4

dBWMR9, dBWMR10, dBWMR11, dBWMR12, dBWMR13, dBWMR14, dBWMR15, dBWMR16, dBWMR17, dBWMR18, dBWMR19, dBWMR20, dBWMR21, dBWMR22, dBWMR23, dBWMR24,

Community 5

vBWMR9, vBWMR10, vBWMR11, vBWMR12, vBWMR13, vBWMR14, vBWMR15, vBWMR16, vBWMR17, vBWMR18, vBWMR19, vBWMR20, vBWMR21, vBWMR22, vBWMR23, vBWMR24,

Community 6

e2VL,

Community 7

CEPDL, URYVR, CEPshVL,

Community 8

vBWMR6,

Community 9

dBWML10,

Community 10

IL2DL, URADL,

Community 11

MCL,

Community 12

dBWML16, dBWML17, dBWML18, dBWML19, vBWML10, vBWML11, vBWML12, vBWML13, vBWML14, vBWML15, vBWML16, vBWML17, vBWML18,

Community 13

dBWML20, dBWML21, dBWML22, dBWML23, dBWML24,

Community 14

IL2DR, URADR,

Community 15

e2D, e2VR, e3D,

Community 16

IL2VR, CEPVR,

Community 17

e3VR,

Community 18

dBWML12,

Community 19

dBWML7, dBWMR7, vBWML4, vBWML5, vBWML6, vBWML7, vBWMR5, vBWMR7, dBWML8, dBWML9, dBWMR8, vBWML8, vBWML9, vBWMR8, exc_gl, hmc,

Community 20

dBWML11,

Community 21

vBWMR4,

Community 22

um2AL, um2AR, um1AL, um1AR, um1PL, um1PR, um2PL, um2PR, vm2AL, vm2AR, vm1AL, vm1AR, vm1PL, vm1PR, vm2PL, vm2PR,

Community 23

SDQL, SDQR, CEPDR, URYDL, OLLL, OLLR, OLQDR, OLQVL, ADAL, RIFR, RIBL, RIBR, RIGL, RIGR, AIBR, RMDDL, RIVL, RIVR, SMDDL, SMDDR, SMDVR, SIBDL, SIBDR, SIBVL, SIBVR, SIADR, DD01, DD02, VD01, VD02, VD03, VD04, VC01, VC02, VC03, VC04,

Community 24

ASER,

Community 25

IL1DL, IL1DR, IL1L, IL1R, IL1VL, IL1VR, RIPR, RMEL, RMER, RMED, RMEV, dBWMR1, dBWMR2, dBWMR3, dBWMR4, dBWMR5, dBWMR6, GLRDL, GLRDR, GLRL, GLRR, GLRVL, GLRVR,

Community 26

ALNR, PQR, ALML, ALMR, AVM, PVM, PLML, PLMR, FLPL, FLPR, DVA, ADER, PDEL, PDER, PHCR, URYDR, AIYL, AIYR, RIS, BDUL, BDUR, PVR, LUAL, LUAR, PVNL, PVNR, AVJL, AVJR, AVDL, AVDR, PVWL, PVWR, RIML, RIMR, AVEL, AVER, RID, AVBL, AVBR, AVAL, AVAR, PVCL, PVCR, RMDR, RMDVR, SABD, SABVL, SABVR, DA01, DA02, DA03, DA04, DA05, DA06, DA07, DA08, DA09, PDA, DB01, DB02, DB03, DB04, DB05, DB06, DB07, AS01, AS02, AS05, AS06, AS10, PDB, VA01, VA02, VA03, VA04, VA05, VA06, VA07, VA09, VA10, VA11, VA12, VB02, VB03, VB04, VB05, VB06, VB07, VB08, VB09, VB10, VB11, HSNL, HSNR,

Community 27

mu_intL, mu_intR, mu_anal, mu_sph,

Community 28

vBWML1, vBWML2, vBWML3,

Community 29

vBWMR1, vBWMR2, vBWMR3,

Community 30

dBWML1, dBWML2, dBWML3, dBWML4, dBWML5, dBWML6, CEPshVR,

Community 31

SAAVR, SMBVL,

Community 32

ASIL, ASIR, AWAL, AWAR, ASGL, ASGR, AWBL, AWBR, ASEL, ADFL, ADFR, AFDL, AFDR, AWCL, AWCR, ASHR, ADLL, ADLR, BAGL, BAGR, URXL, URXR, ADEL, IL2L, IL2R, IL2VL, CEPVL, OLQDL, OLQVR, AINL, AINR, AIML, RIH, URBL, URBR, RIR, AIAL, AIAR, AUAL, AUAR, AIZL, AIZR, ADAR, RMGL, RMGR, AIBL, RICR, SAADR, SAAVL, RIAL, RIAR, RMFL, RIPL, URAVL, RMDDR, RMDL, RMDVL, RMHL, RMHR, SMDVL, SMBVR, SIAVL, SIAVR, VB01,

Community 33

vBWML19, vBWML20, vBWML21, vBWML22, vBWML23,

Community 34

DD03, DD04, VD05, VD06, VD07, VD08, VD09,

Community 35

URYVL, RIFL, URAVR, AS04, VA08,

Community 36

I1L, I1R, I2L, I2R, I3, I4, I5, I6, M1, M2L, M2R, M3L, M3R, M4, M5, MCR, MI, NSML, NSMR, pm1, pm2D, pm2VL, pm2VR, pm3D, pm3VL, pm3VR, pm4D, pm4VL, pm4VR, pm5D, pm5VL, pm5VR, pm6D, pm6VL, pm6VR, pm7D, pm7VL, pm7VR, pm8, mc1DL, mc1DR, mc1V, mc2DL, mc2DR, mc2V, mc3DL, mc3DR, mc3V, e3VL, g1AL, g1AR,
