## Supplementary figures and images for "Functions of *C. elegans* neurons from synaptic connectivity"

### Supplementary File 5

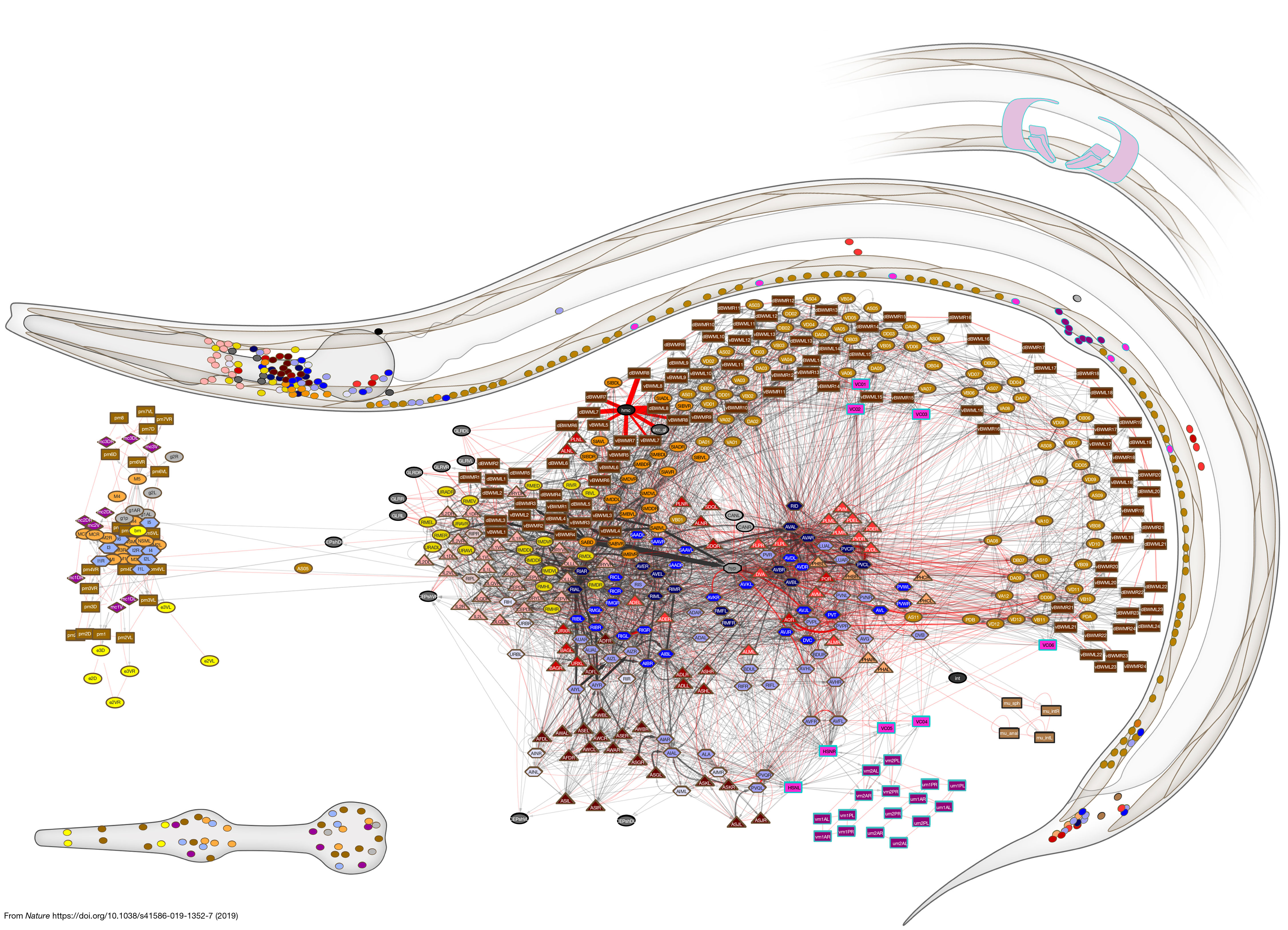

### Supplementary File 7

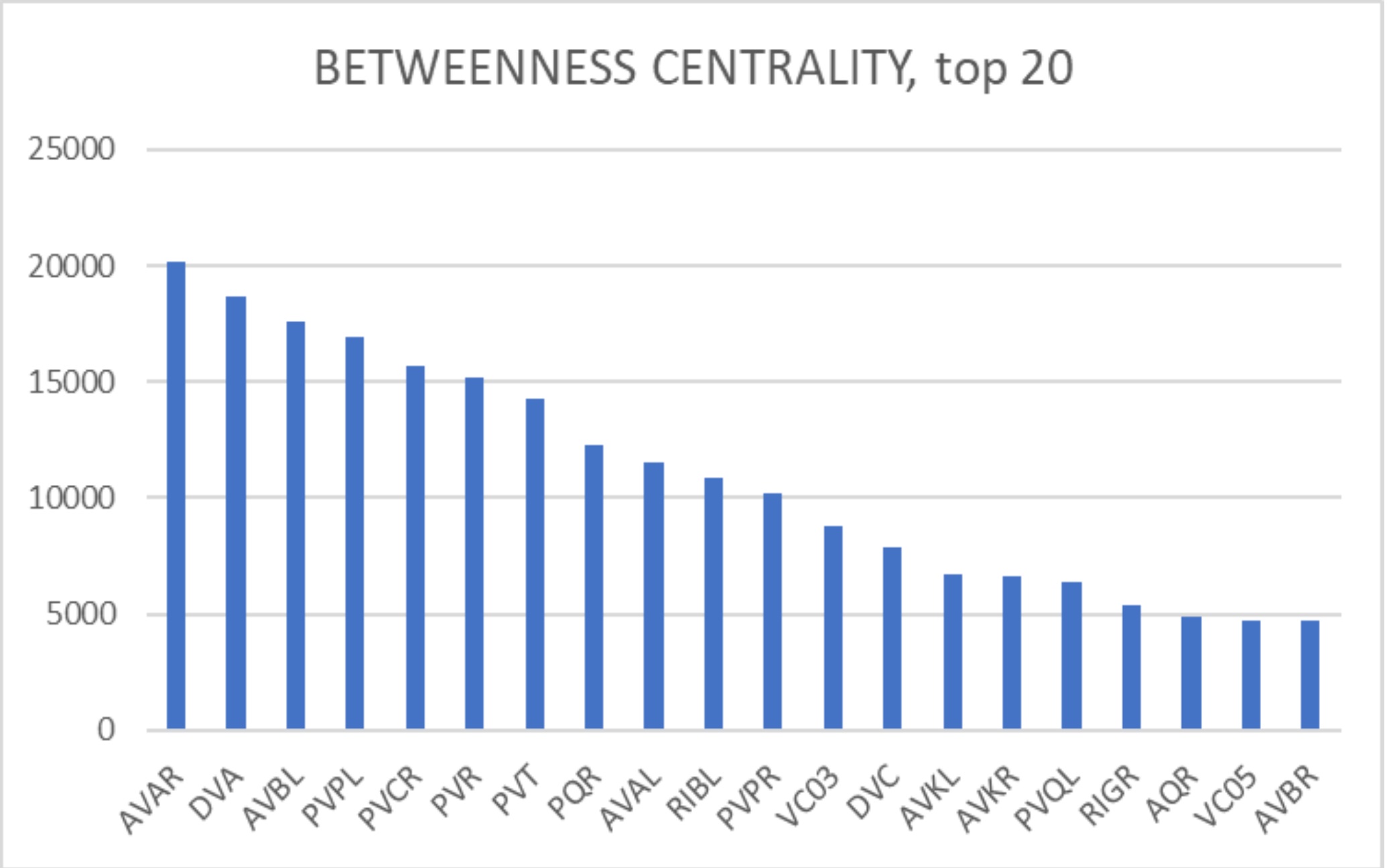
